# Discovery of Competent Chromatin Regions in Human Embryonic Stem Cells

**DOI:** 10.1101/2023.06.14.544990

**Authors:** Julian Pulecio, Zakieh Tayyebi, Dingyu Liu, Wilfred Wong, Renhe Luo, Jeyaram R Damodaran, Samuel Kaplan, Hyunwoo Cho, Jielin Yan, Dylan Murphy, Robert W. Rickert, Abhijit Shukla, Aaron Zhong, Federico González, Dexin Yang, Wenbo Li, Ting Zhou, Effie Apostolou, Christina S. Leslie, Danwei Huangfu

## Abstract

The mechanisms underlying the ability of embryonic stem cells (ESCs) to rapidly activate lineage-specific genes during differentiation remain largely unknown. Through multiple CRISPR-activation screens, we discovered human ESCs have pre-established transcriptionally competent chromatin regions (CCRs) that support lineage-specific gene expression at levels comparable to differentiated cells. CCRs reside in the same topological domains as their target genes. They lack typical enhancer-associated histone modifications but show enriched occupancy of pluripotent transcription factors, DNA demethylation factors, and histone deacetylases. TET1 and QSER1 protect CCRs from excessive DNA methylation, while HDAC1 family members prevent premature activation. This “push and pull” feature resembles bivalent domains at developmental gene promoters but involves distinct molecular mechanisms. Our study provides new insights into pluripotency regulation and cellular plasticity in development and disease.

**One sentence summary:** We report a class of distal regulatory regions distinct from enhancers that confer human embryonic stem cells with the competence to rapidly activate the expression of lineage-specific genes.

## INTRODUCTION

Embryonic stem cells (ESCs) have unique chromatin features that are thought to provide the molecular basis for their developmental plasticity (*1–3*). The presence of bivalent domains at the promoters of lineage genes as well as the hyperdynamic binding of chromatin-associated proteins have been suggested to support the ability of ESCs to swiftly activate or repress distinct lineage-specific gene expression programs in response to external differentiation cues. However, it remains unclear how ESCs can rapidly establish distal enhancer elements to activate the expression of lineage-specific genes upon differentiation. Previous studies have suggested that chromatin regions that become active enhancers in differentiated cells may already be premarked in ESCs. For example, early studies have shown that several enhancers controlling the expression of blood or hepatocyte specific genes are marked with unmethylated CpGs in mouse ESCs (*4, 5*). More recent works examined the temporal dynamics of multiple chromatin features during neural and endodermal lineage differentiation. They observed that the regions identified as active enhancers by the presence of H3K27ac in differentiated cells tended to have certain chromatin marks prior to differentiation. Some enhancer regions are premarked in ESCs by the histone marks H3K27me3 (*6*) and H3K9me3 (*7*) and the histone variant H2A.Z (*8*), before acquiring the primed enhancer mark H3K4me1 and eventually becoming active enhancers marked by H3K27ac (*9, 10*). These findings suggest that some tissue-specific enhancers are premarked in ESCs with permissive features that could support their competence to induce gene expression in response to the binding of lineage-specific transcription factors (TFs). However, functional assays that do not rely on inferred correlations are needed to identify specific regions with the competence to activate gene expression upon TF binding in ESCs. Such investigations could help us to better define specific premarking features and understand how they contribute to the transcriptional competence of the chromatin. Addressing these questions is critical for understanding pluripotency regulation in ESCs, the acquisition of pluripotency during somatic cell reprogramming and more broadly, the molecular mechanisms that sustain cellular plasticity in development, regeneration, and pathological conditions such as cancer.

We set out to establish a functional assay to examine the presence of distal (non-promoter) chromatin regions with the competence to activate the expression of lineage-specific genes in human ESCs (hESCs). We postulated that the dead Cas9 (dCas9) protein could function as a pioneer TF (*11*), by binding to previously inaccessible chromatin regions through RNA-guided targeting (*12*) but without the constraint of TF binding motifs. In addition to opening the chromatin, the interrogation of competence also requires the recruitment of transcriptional activators, for which we employed the CRISPR activation (CRISPRa) systems previously shown to induce gene expression when targeted to gene promoters or active enhancers (*13–16*). Employing CRISPRa screens to systematically interrogate five lineage-specific gene loci that were inactive at the ESC stage, we discovered defined chromatin regions in hESCs, which we termed Competent Chromatin Regions (CCRs), that are capable of inducing gene expression over long distances upon targeting with strong transcriptional activators. To identify features that could distinguish CCRs from non-CCRs (regions not capable of responding to CRISPRa), we conducted a comprehensive analysis encompassing three key aspects of local chromatin state: three-dimensional (3D) chromatin conformation, DNA sequence features, and occupancy of chromatin factors at the interrogated loci. We found that CCRs are exclusively localized within the same topologically associated domains (TADs) as their target genes. They can be located in distal regions far from their target gene, and interestingly, many of them do not show enriched chromatin contacts with their associated genes. Unexpectedly, CCRs cannot be distinguished from non-CCRs based on the enrichment of the histone features previously associated with the premarking or priming of enhancers. Instead, CCRs showed a significant enrichment of the POU sequence motifs and displayed higher levels of binding by pluripotency TFs OCT4 (POU5F1) and NANOG when compared to non-CCRs.

Furthermore, CCRs are characterized by the enriched presence of activating chromatin factors, such as DNA demethylation factors TET1 and QSER1, and repressive factors like histone deacetylase (HDAC) enzymes. This resembles the opposing features found in bivalent domains associated with lineage gene promoters. We further demonstrate that the HDAC enzymes restrict the transcriptional competence of CCRs while QSER1 and TET1 protect it by inhibiting the hypermethylation of the DNA. In the context of the early stages of ESC differentiation, these chromatin features could facilitate the binding of pioneer TFs such as FOXA2, thus readily supporting the activation of lineage-specific gene expression. This study advances our understanding of pluripotency regulation and developmental plasticity by identifying a distinct type of distal regulatory region that enable ESCs to activate lineage gene expression independently of the establishment of comprehensive, lineage-specific gene regulatory programs.

## RESULTS

### A CRISPRa screen discovers CCRs in hESCs

We hypothesize that hESCs have distal chromatin regions capable of activating lineage-specific genes when targeted by CRISPRa. To test this, we began with a comprehensive case study on *PDX1*, a lineage-specific gene that is inactive at the ESC stage but is expressed in pancreatic progenitor (PP) cells (*17*). We utilized our established H1 PDX1^GFP/+^ line (*18*) to monitor *PDX1* expression and derived two hESC lines for inducible CRISPRa targeting. The first cell line was engineered for doxycycline-inducible expression of the targetable activator dCas9-VP64, and the second line included both dCas9-VP64 and the MS2-p65-HSF1 complex to form the inducible Synergistic Activation Mediator (iSAM) system (*19*) (Supp Fig. 1a). Next, we introduced a lentiviral library of ∼7,500 gRNAs spanning a 100kb region surrounding the *PDX1* locus (Supp Fig. 1c). After treating the transduced cells with doxycycline for four days in the E8 hESC maintenance media, flow cytometric analysis was performed to assess *PDX1-*GFP expression. The iSAM system outperformed the dCas9-VP64 system and generated a substantial PDX1-GFP+ cell population (Supp Fig. 1d). Therefore, we sorted GFP+ and GFP-cells generated through the iSAM system and assessed the relative abundance of the gRNAs in each cell population by next generation sequencing (NGS). After excluding gRNAs with low counts, we identified that the positive control gRNAs targeting the *PDX1* promoter region were enriched in the PDX1-GFP+ cell population (compared to the PDX1-GFP-) with an average log2 fold change (LFC) = 1.05 while the negative control gRNAs had a LFC = -0.04. We used the average LFC of the positive control gRNAs as a threshold and detected 423 gRNAs enriched in the PDX1-GFP+ fraction with a LFC >1 that are not targeting *PDX1* gene body or promoter regions (Fig. 1a, Supp. Fig. 1e, Supp Table S1). To better define the chromatin regions associated to the gRNAs enriched in GFP+ cells, we performed MAGeCK analysis and applied a sliding window approach (*20, 21*) that allowed us to specify six Competent Chromatin Regions (CCRs) defined by their ability to induce *PDX1* expression in hESCs when targeted by the CRISPRa iSAM tool (IDR < 0.05, |Z-score| > 0.25, average size = 265bp, see methods) (Fig. 1c, Table S2). We observed that some CCRs were located as far as 20 kb from the transcription start site (TSS) of *PDX1*, whereas other regions in closer proximity to the *PDX1* TSS did not respond to CRISPRa interrogation. These findings indicate that the ability to respond to CRISPRa is not uniform across all regions surrounding a target gene, and that factors beyond the linear distance from the associated gene may determine the transcriptional competence of a chromatin region.

**Figure 1.**
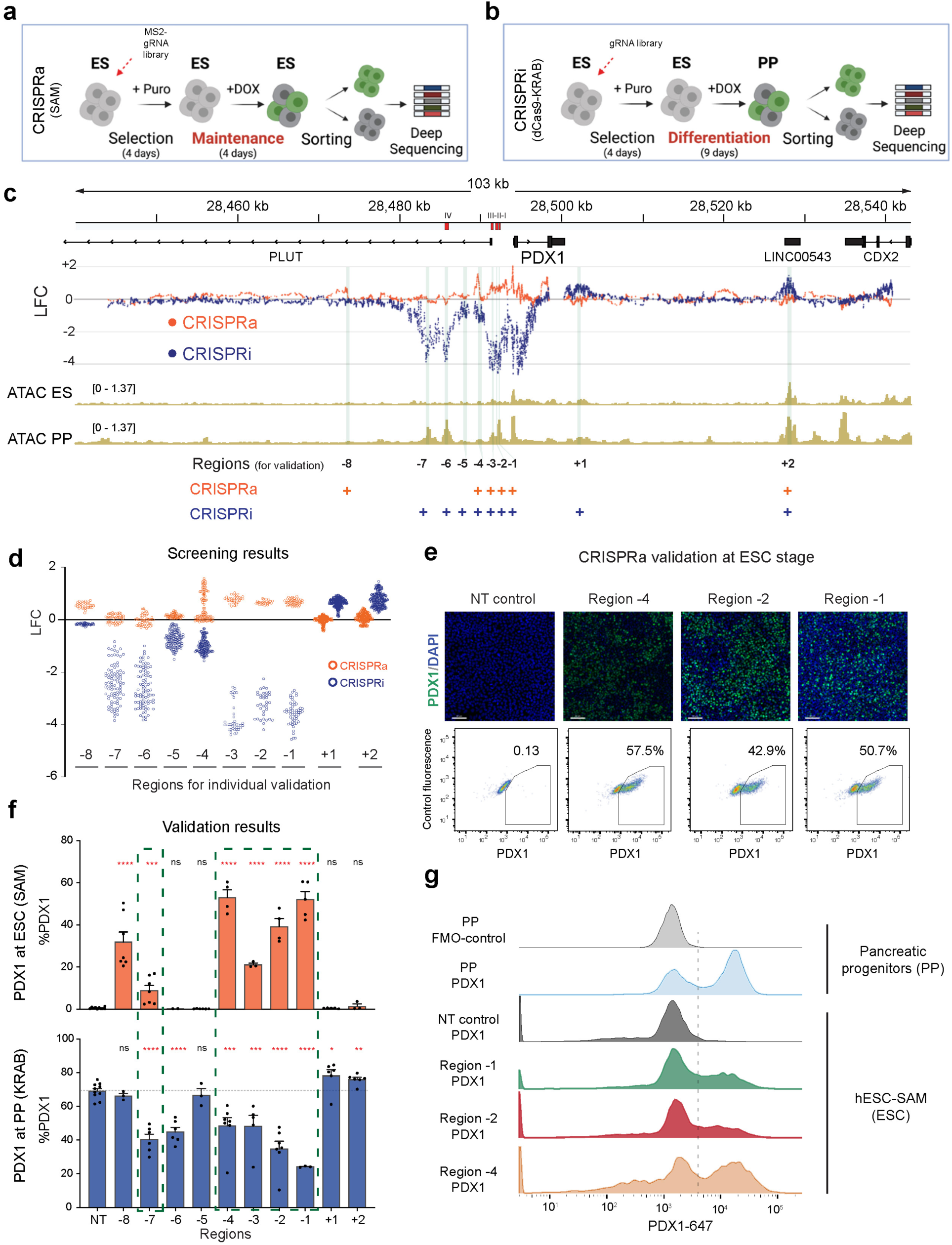
CRISPRa-based interrogation to discover transcriptionally competent regions in hESC. **a)** Schematics of the CRISPR-a screen designed to discover transcriptionally competent regions to promote PDX1 expression at the ESC stage. **b)** CRISPR-i screen designed to discover PDX1 enhancers required during the pancreatic lineage-specification. **c)** Results of the CRISPR-based screens (CRISPRa: orange, CRISPRi: blue), each dot depicts the enrichment of a gRNA in the PDX1+ (positive LFC) or PDX1- (negative LFC) fractions. As a reference, previous candidate PDX1 enhancers (Area I-IV) are indicated as red bars on top and ATAC-seq tracks at the ESC and PP stages are included. Regions discovered by each screen using a sliding window analysis and selected for individual validation are depicted in the lower part and highlighted in green (+ indicates the CRISPR screen used to discover the candidate regulatory region). **d)** Magnification of the candidate PDX1 regulatory regions used for individual validation. **e)** Immunostaining images and FACS dot plots showing PDX1 expression levels after interrogation of a subset of candidate regulatory regions with the SAM tool at the ESC stage. A non-targeting gRNA was used as a control (see Methods) (PDX1 = Green, DAPI = blue). **f)** Summary of the flow cytometry experiments to validate the CCRs that are able to control PDX1 expression at the ESC stage (CRISPRa: orange) and the enhancers required to express PDX1 during the pancreatic differentiation (CRISPRi: blue). The green dashed line indicates the CCRs that overlap with PDX1 enhancers (n ≥ 3). **g)** Representative histograms comparing the PDX1 protein expression levels in differentiated Pancreatic Progenitors and hESC-SAM targeting three CCRs (A Fluorescence Minus One (FMO) control for the PP cells and a non-targeting gRNA control for the hESC-SAM are included).

### CCRs and tissue-specific enhancers have substantial overlaps

To identify factors that could contribute to the transcriptional competence of CCRs, we first evaluated whether these regions exhibited enhancer-associated chromatin features. In undifferentiated ESCs, the six CCR hits had weak or no detectable ATAC, H3K4me1 or H3K27ac signals. However, upon differentiation into PP cells (*22*), these CCR hits exhibited elevated levels of these signals (Fig. 1c, Supp Fig. 1f). Therefore, CCRs do not constitute primed or active enhancers at the ESC stage, but they appear to overlap with the genomic regions that acquire enhancer activities after lineage specification. To systematically compare CCR hits with functional *PDX1* enhancers, we performed a CRISPRi screen aimed at identifying regulatory regions necessary for *PDX1* expression during pancreatic differentiation. First, we generated a doxycycline-inducible dCas9-KRAB line using the H1 PDX1^GFP/+^ line (*18*)(Supp. Fig1b). Next, we transduced the cells with a tiled gRNA library targeting the *PDX1* locus, matching the gRNAs used in the CRISPRa screen but adapted for the dCas9-KRAB system. Transduced cells were treated with doxycycline to induce dCas9-KRAB expression during pancreatic differentiation (Fig. 1b). After nine days of differentiation, GFP+ and GFP-fractions were enriched by fluorescence-activated cell sorting (FACS), and gRNA abundance was determined by NGS (Supp Fig. 1e, Table S1). We identified 840 gRNAs significantly enriched in either fraction (|LFC| > 1), and MAGeCK and sliding window analyses revealed twelve *PDX1* regulatory region hits (IDR < 0.001, |Z-score| > 3, average size = 949bp, see Methods) (Fig. 1c, Table S2). Some hits overlap with sequences that are homologous to known *Pdx1* enhancers in mice (*23, 24*) (Table S2), demonstrating functional conservation of these enhancers across species.

The CRISPRa and CRISPRi screens identified several overlapping regions as well as regions that were unique to each screen. To account for potential false negatives and positives, we selected 10 regions for individual validations, including 5 regions identified by both screens and 5 regions exclusive to either screen (Fig. 1c-d, Table S2). Following the approach of the screens, we performed the CRISPRa validation in undifferentiated hESCs to examine the sufficiency of a region to activate *PDX1* expression and conducted the CRISPRi validation in cells undergoing pancreatic differentiation to evaluate any effects on *PDX1* induction. The CRISPRi validation experiments revealed 6 enhancers that when targeted led to reduced PDX1 expression and 2 regions that caused increased PDX1 expression compared to non-targeting (NT) gRNA controls based on flow cytometric analysis and immunostaining (Fig. 1f, Supp Fig. 1g). Meanwhile, the CRISPRa experiments identified 6 regions that were competent to activate PDX1 expression in undifferentiated hESCs, with 5 of these regions overlapping with the enhancers validated by CRISPRi (Fig. 1e-f). Remarkably, CRISPRa targeting of a single CCR can induce the expression of PDX1 in hESCs at comparable protein levels to those observed in PP cells derived from the guided differentiation of hESCs where PDX1 expression is supported by a multilayered gene regulatory network (Fig. 1g). Finally, to further validate our results, we generated several CRISPR-Cas9 edited lines for two of the hits identified by both CRISPRa and CRISPRi (*PDX1* Region-7 (R-7) and *PDX1* Region-1 (R-1)) and confirmed that the deletion of either region caused a significant reduction of *PDX1* expression during pancreatic differentiation (Supp Fig. 1h). Together, our results demonstrate that the CRISPRa system can be employed as a targetable pioneer TF and activator to identify CCRs capable of activating the expression of lineage-specific genes, such as *PDX1*, without having to expose hESCs to any differentiation signals. Moreover, CCRs and tissue-specific enhancers have substantial overlaps, indicating that many chromatin regions associated with lineage-specific enhancers are already competent to support transcription in undifferentiated ESCs, prior to the establishment of lineage-specification programs.

### CCRs are present across extended genomic distances in multiple developmental gene loci

To determine the generalizability of our findings at the *PDX1* locus and to better characterize the molecular features of the CCRs, we broadened our search to encompass four additional developmental genes that become activated during the differentiation of hESCs to the definitive endoderm (DE) lineage. Since our findings at the *PDX1* locus suggest that most CCRs later become enhancers in differentiated cells, we focused on regions (excluding promoters) that are accessible at either the ESC or DE stage based on our ATAC-seq data (*25*). This targeted interrogation strategy allowed us to simultaneously expand our screen to more genes and span greater genomic distances. We employed a gRNA library including ∼5,400 gRNAs that targeted 163 regions encompassing a total of 12.8 Mb distance surrounding the *GATA6*, *EOMES*, *SOX17* and *MIXL1* loci. After transduction of the gRNA library into the iSAM hESC line, cells were treated with doxycycline for four days in the E8 hESC maintenance media. Next, intracellular staining was performed for GATA6, EOMES, SOX17 or MIXL1 followed by FACS sorting (Fig. 2a, Supp Fig. 2a), and all positive fractions and all negative fractions were combined for NGS. After filtering out potential off-target gRNAs, we identified 197 hits that were significantly enriched in the combined positive fractions (Table S3, Fig. 2b-d, Supp. Fig. 2b-d). The genomic regions that contained at least two significantly enriched gRNAs within a 100bp range were considered CCR hits (average size = 355bp) (see methods). This strategy allowed us to identify CCRs linked to the activation of GATA6 (16 hits), *SOX17* (8), *EOMES* (7) and *MIXL1* (5). We selected 15 CCR hits for individual validations using gRNAs targeting the CCRs coupled with iSAM activation at the ESC stage, and all 15 hits were able to induce the expression of their corresponding genes as determined by flow cytometry and immunofluorescence assays (Fig. 2e-f, Supp. Fig. 2f). Similar to the *PDX1* experiments, targeting a single CCR in undifferentiated hESCs was sufficient to activate the expression of the target genes to comparable levels as those observed in differentiated cells (Supp. Fig. 2g). Therefore, the CCRs are highly potent and capable of driving lineage-specific gene expression, which is normally achieved during development through a complex regulatory cascade that orchestrates gene expression programs.

**Figure 2.**
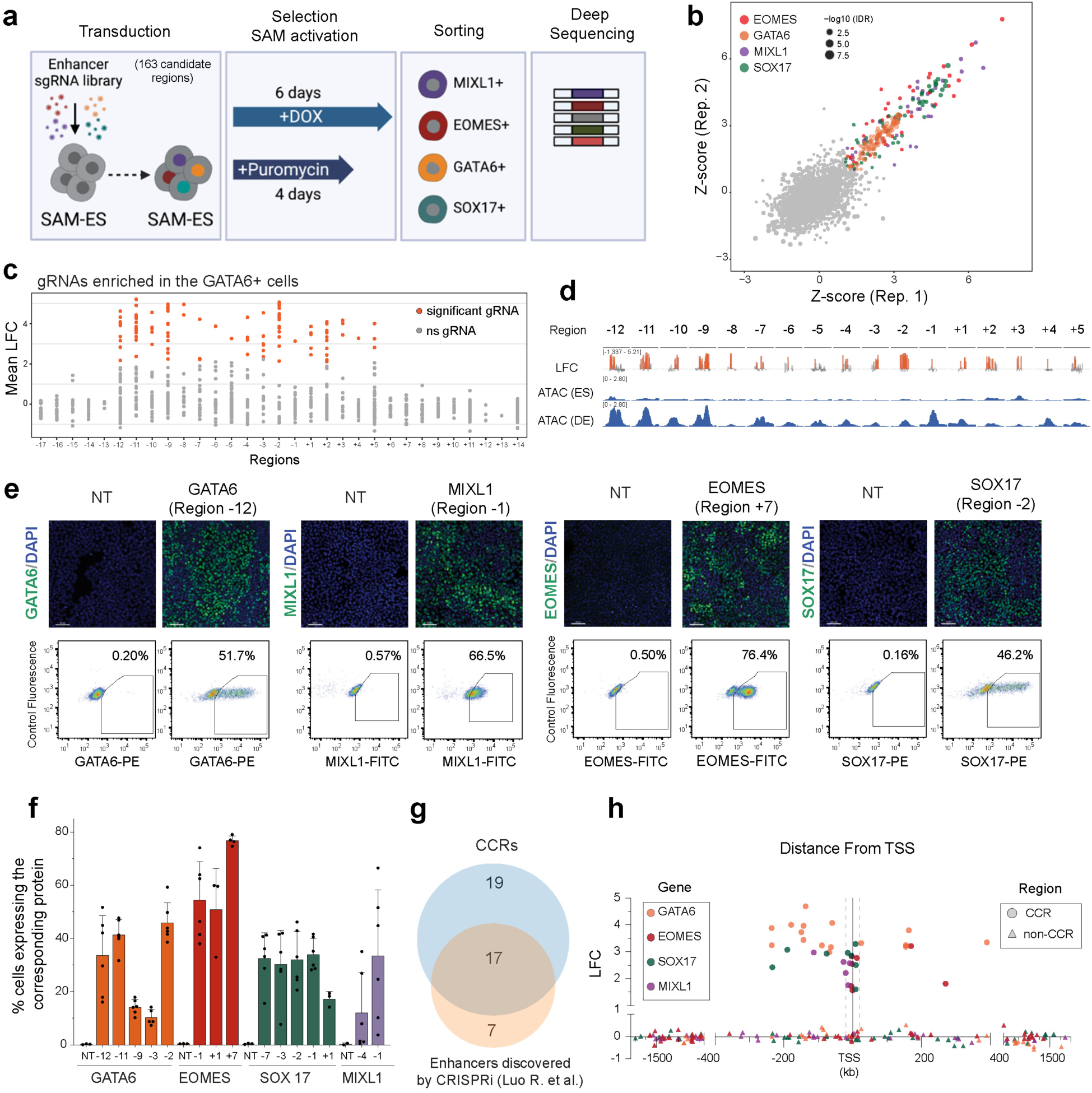
Broad interrogation for CCRs able to activate the expression of multiple developmental genes in hESCs. **a)** A tiled gRNA library targeting differential ATAC peaks detected during the transition from ESC to DE coupled with the depicted CRISPRa screening strategy was used to discover the CCRs of the genes GATA6, EOMES, SOX17 and MIXL1 at the ESC stage. **b)** Reproducibility between replicates of the screens performed with the iSAM-hESC line at the ESC stage based on the Z-score (see methods). Each dot represents a gRNA inside the interrogated regions, color-coded for the associated gene and size-coded for IDR (-log10) (ns = non-significant). **c)** LFC distribution (GATA6+/GATA6-) of the gRNAs targeting the 31 regions interrogated around the GATA6 locus after the CRISPRa screen. Each dot represents a gRNA color-coded as significantly (orange) or non-significantly enriched (gray) in the FACS-sorted positive fraction (see methods). **d)** Multi-track plot amplification for each interrogated region surrounding the GATA6 locus containing at least one sgRNA significantly enriched in the positive fraction (each bar in the LFC track corresponds to a gRNA and is color-coded as in 2c). ATAC-seq signal at ESC and DE stages are included. **e)** Representative immunofluorescence images (*upper panel*) and FACS plots (*lower panel*) to show the effect of activating individual CCRs via gRNA-targeting with the SAM tool for each gene, compared to non-targeting (NT) gRNA controls (scale bar =80um). **f)** Summary of the validation experiments performed in multiple candidate CCRs for each gene. Each bar corresponds to the percentage of positive cells expressing the protein associated to each model gene after CCR interrogation via CRISPRa targeting (n >4, NT= cells transduced with non-targeting gRNA controls). **g)** Intersection between functionally validated enhancers and the discovered CCRs of the four interrogated genes (table s3). **h)** Genomic location and LFC of each CCR and non-CCR based on the distance from the TSS of their associated gene. Each region is shape-coded by its chromatin competence and color-coded by its associated gene.

Next, we analyzed the dynamics of the chromatin features commonly associated to enhancer activity in CCRs during the transition from ESC to DE cells. Our data revealed that CCRs in ESCs typically exhibited low levels of ATAC, H3K4me1 and H3K27ac signals. However, when ESCs were differentiated to the DE stage, these CCRs became more accessible and showed increased levels of H3K4me1 and H3K27ac (Fig. 2d, Supp. Fig. 2e, 2h). We further compared the same chromatin features between CCRs and non-CCRs (regions that did not cause any activation of their linked genes upon CRISPRa interrogation) and found that H3K27ac levels were significantly higher in CCRs at the DE stage, strongly suggesting that CCRs tend to become active enhancers in differentiated cells (Supp. Fig. 2i-k). To validate this observation we directly compared the genomic regions covered by CCRs and CRISPRi validated enhancers of the same four DE genes recently discovered by our group (*26*), and we found a substantial overlap between CCRs in hESCs and developmental enhancers in differentiated cells (Fig. 2g, Table S3). The CRISPRa screen on DE genes targeted more genomic loci and covered greater distances compared to the one conducted on *PDX1*, enabling us to examine the distribution of the CCRs more comprehensively across genomic locations relative to the target gene. CCRs spanned variable genomic distances relative to their associated genes, with ∼38% of them located more than 100 kb away from the TSS of the corresponding gene (Fig. 2h). Together, these results reveal a strong overlap between CCRs and enhancers, and indicate that CCRs, like enhancer elements, are not restricted to the immediate linear vicinity of their associated genes. Our findings strongly support the notion that the pre-establishment of transcriptionally competent regions prepares ESCs for rapid activation of lineage-specific gene expression programs upon exposure to differentiation signals.

### CCRs are in the same topologically associated domains as their corresponding genes

With an expanded set of 42 CCRs discovered across five developmental gene loci (Table S3), we further investigated the genomic characteristics that distinguish CCRs from non-CCRs. Considering that CCRs were not found at chromatin regions with clear H3K27ac and H3K4me1 enhancer marks at the ESC stage, we explored other features associated with enhancer functionality. Chromatin conformation assays have suggested that gene regulatory regions, including enhancers, are typically located within topologically associated domains (TADs) containing their target genes (*27*). TADs are thought to facilitate promoter-enhancer interactions either through direct contact or by oligomerization of the TFs bound to these regions (*28*). To determine whether CCRs are also confined within TADs, we performed Hi-C assays on undifferentiated hESCs and used TopDom (*29*) to identify TADs, defined as regions with larger contact frequencies within than outside of the domain. All the genes except for *MIXL1* were assigned to a TAD, and we found that their respective CCRs were exclusively located within the same TAD (Fig. 3a-b). In contrast, ∼59% of the non-CCRs were found outside the TADs linked to the corresponding genes (Fig. 3a-b). Considering the choice of window size used to scan for TAD boundaries can affect the TAD assignment, we expanded the window size from 5 to 10 bins and identified a *MIXL1*-containing TAD (Supp. Fig. 3d). All five *MIXL1* CCRs were found within this TAD, whereas ∼74% *MIXL1* non-CCRs were outside of it (Table S4). Further examination of CCRs and the TADs across all five loci showed a concordant asymmetric distribution of CCRs where the TAD is located on one side of the TSS, such as in the case of *EOMES* and *SOX17*. Specifically, CCRs were located on one side of the TSS, within the TAD, while the regions on the other side were non-CCRs, despite no difference in their distance to the TSS. These findings suggest that the three-dimensional organization of the chromatin is a crucial spatial constraint of CCRs.

**Figure 3.**
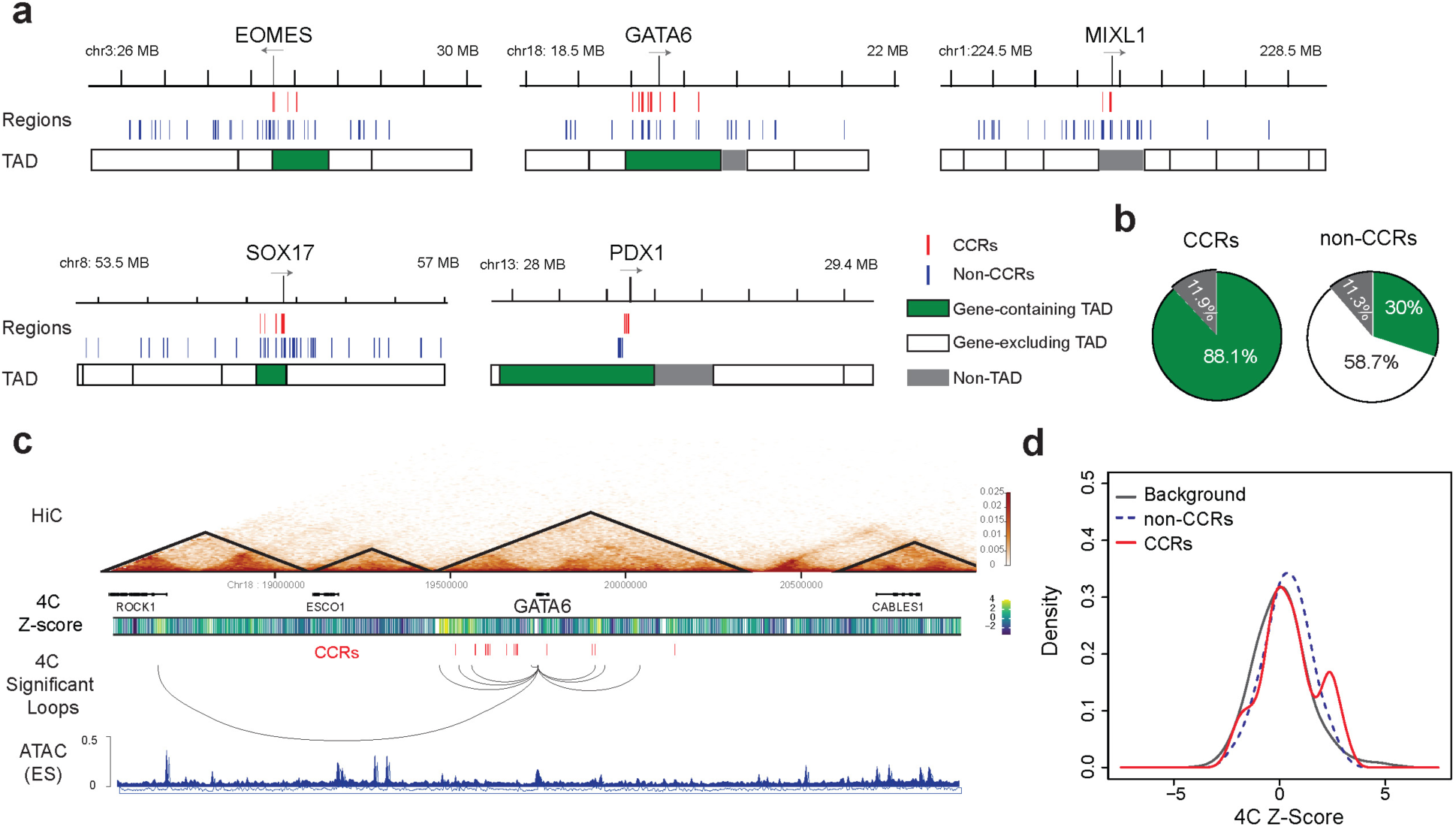
Chromatin conformation at the CCRs containing loci. **a)** TAD distribution revealed by Hi-C assays around the five interrogated genes at the ESC stage. Gene strand direction is indicated (grey arrows), CCRs are depicted by red bars and non-CCRs by blue bars. **b)** Percentage of CCRs and non-CCRs located inside the TADs containing their associated genes. **c)** Hi-C heatmap using KR normalization to visualize chromatin contacts surrounding the GATA6 locus at the ESC stage (50kb resolution). TADs are depicted as black lines. (*Mid panel*) 4C experiments using as a viewpoint the GATA6 gene at the ESC stage (n=3). A heatmap based on the Z-score is shown to describe the local intensity of the signal using a sliding windows strategy (see methods). (*Low panel*) Significant contacts (q ≤ 0.1) in the 4C data. GATA6 CCRs are depicted as red bars. As a reference ATAC signal at the ESC stage is included. **d)** 4C Z-scores for contacts across the GATA6, SOX17, PDX1, and EOMES experiments were used for kernel density estimates across different types of CCRs. The background regions were defined as regions that did not overlap competent or non-competent regions.

To further investigate the spatial relationship of CCRs and their corresponding genes, we next explored whether CCRs already have higher levels of chromatin interactions with the corresponding genes in hESCs compared to non-CCRs, before any activation induced by CRISPRa targeting or cellular differentiation could take place. Due to the limited resolution of the Hi-C data for identifying chromatin loops in bins smaller than 50kb, we performed 4C-seq (4C) assays on hESCs using *EOMES*, *PDX1*, *SOX17* and *GATA6* as viewpoints (VPs) (Table S5). To determine the chromatin contacts with the VPs we computed kernel density estimates of the 4C Z-Scores for CCRs, non-CCRs and the remaining regions, by calculating the probability of observing the counts under an estimated background model (Methods). The non-CCRs exhibited a unimodal density, suggesting that the non-competent regions are neither preferentially enriched nor depleted of contacts with the corresponding gene (Fig. 3c-d, Supp. Fig. 3a-c,e). In comparison, CCRs tended to have a higher number of significant chromatin contacts with their corresponding VPs compared to background levels, but only a subset of CCRs showed a strong contact enrichment with the corresponding genes as indicated by the bimodal density estimate (Fig. 3d). Interestingly, CCRs found at distant locations tended to have a higher number of chromatin contacts with their associated genes (Supp. Fig. 3f). In summary, our Hi-C and 4C results demonstrate that CCRs are confined within the same TADs of their corresponding genes, suggesting that their activity is constrained by the three-dimensional conformation of the chromatin. In addition, while 3D spatial proximity to the target gene may contribute to CCR function, it is not an absolute requirement.

### CCRs are preferentially bound by pluripotent TFs and chromatin activators

To investigate if additional molecular factors beyond chromatin conformation could influence the transcriptional competence of a chromatin region in ESCs, we first examined DNA sequence features. Enhancers contain DNA motifs that allow the sequential binding of lineage-specifying master TFs and other co-factors to recruit the transcriptional machinery (*30*). Since CCRs are identified before the expression of endogenous lineage-specifying master TFs, we hypothesized that CCRs may contain DNA sequence features that recruit factors other than the master TFs to establish the transcriptional competence. To test this, we performed HOMER motif enrichment analysis (*31*) comparing CCRs to random genomic regions and discovered that CCRs are significantly enriched in POU, HMG(SOX), ZF (GATA) and FOX domains (Fig. 4a). Furthermore, motif enrichment analysis comparing CCRs and non-CCRs within the TADs containing our model genes revealed a significant enrichment of POU motifs in CCRs (Fig. 4b, Supp. Fig. 4b). This enrichment of POU motifs was consistently observed when comparing CCRs to all non-CCRs, irrespective of their location relative to the TAD (Supp. Fig. 4a, Table s6).

**Figure 4.**
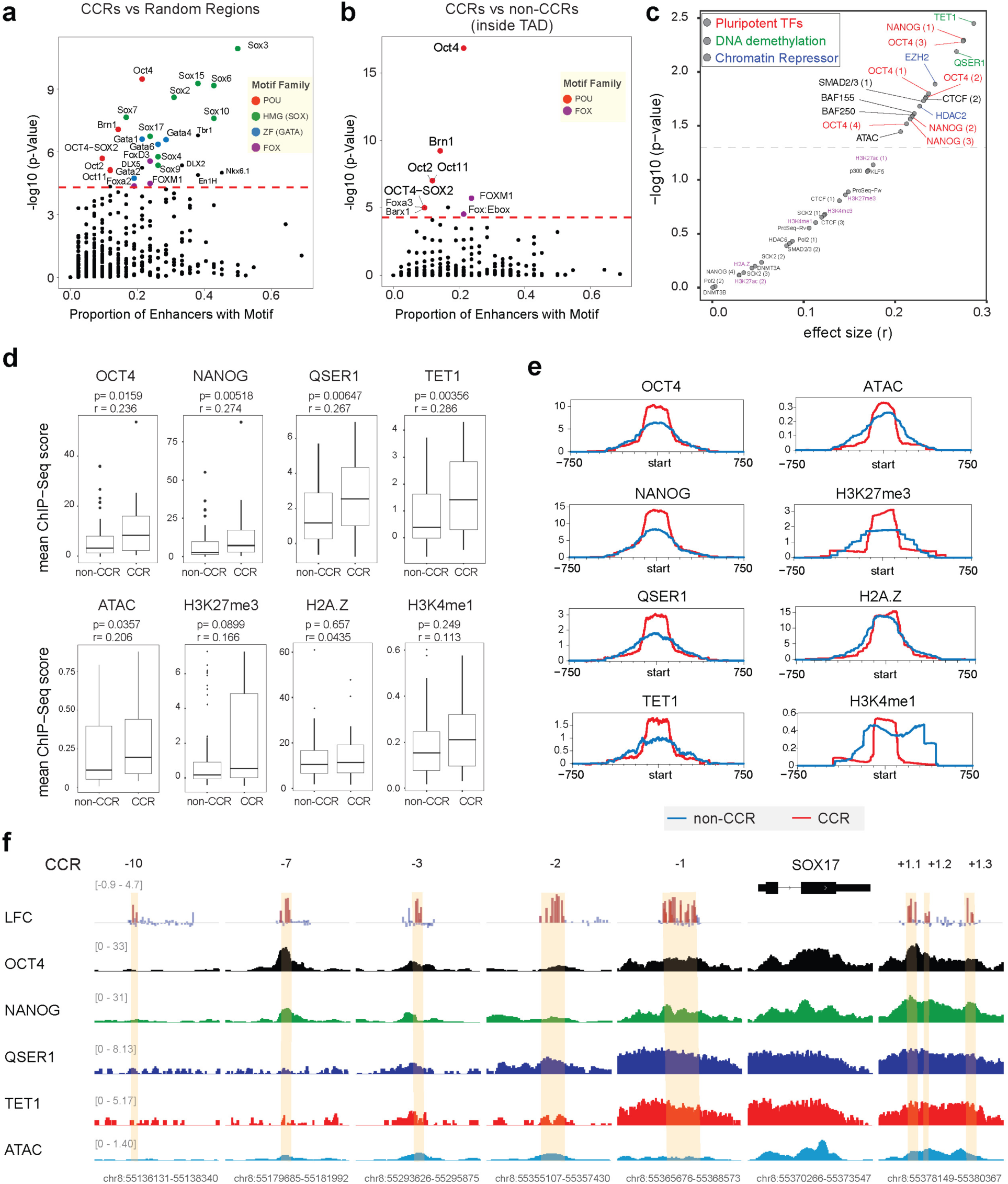
Analyses of the DNA features and chromatin-associated factors enriched at the CCRs. **a)** HOMER motif enrichment analysis plot to summarize the significantly enriched TF motifs (above the red-dotted line) when comparing CCRs (42) vs random genomic regions (48144). **b)** HOMER motif enrichment analysis plot to summarize the significantly enriched TF motifs (above the red-dotted line) when comparing CCRs vs non-CCRs inside the same TAD (62). **c)** ChIP-seq data collected at the ESC stage was used to measure the signal levels of each chromatin-associated factors in the CCRs vs non-CCRs (inside the gene-containing TADs). Factors that bind significantly differently between CCRs and non-CCRs are indicated above the blue-dotted line (Wilcoxon rank-sum test, p< 0.05, effect size >0.2). Categories of chromatin-associated factors enriched in the CCRs are color-coded. Histone marks previously associated to enhancers are highlighted in purple. **d)** Average ChIP-seq scores of chromatin features measured at CCRs and non-CCRs in hESCs (Wilcoxon rank-sum test, p values and effect sizes indicated on top). **e)** Average of normalized counts visualized as a MetaPeak for the main ChIP-signal data sets enriched in CCRs (red), compared to the non-competent regions(blue). **f**) Multi-track plot to illustrate the ChIPseq signal of the enriched factors at the CCRs around the SOX17 locus. CCRs are highlighted in yellow and red bars depict the log Fold Change (LFC) in the CRISPRa screen for individual gRNAs inside each CCR (blue bars = gRNAs not enriched in CCRs).

The enrichment in POU motifs at the CCRs suggests an increased presence of OCT4 and related pluripotency factors in these regions. We compared the binding levels of OCT4 in hESCs between CCRs and non-CCRs within gene-containing TADs, along with 24 common chromatin-associated factors including pluripotency TFs, histone marks and chromatin modifiers (Table S7). Our analyses revealed that among various hits, the top factors enriched at CCRs were the pluripotent TFs OCT4 and NANOG (enriched in multiple hESC lines) and the DNA demethylation factors QSER1 and TET1 (*32*) (Wilcoxon rank-sum test, r >0.2, p <0.05) (Fig. 4c, Table S7). In contrast, CCRs did not show an increased presence of the histone modifications H3K4me1 and H3K27ac, histone marks commonly associated to primed and active enhancers. Additionally, levels of the histone modification H3K27me3 or histone variant H2A.Z, previously associated to poised enhancers or “pre-enhancers” in ESCs, were not significantly different between CCRs and non-CCRs (Fig. 4d) (*6, 8*). However, we did observe a modest but statistically significant increase in chromatin accessibility in CCRs compared to non-CCRs inside the same TADs (Fig. 4d). Visual inspection of the ChIP-seq profiles together with meta-signal plots confirmed that the binding levels of most of the enriched factors in CCRs were indeed higher than those in non-CCRs inside gene-containing TADs (Fig. 4e-f, Supp. Fig. 4d-e), suggesting that the increased binding of these factors may contribute to transcriptional competence. A comparison between CCRs and non-CCRs without the TAD restriction showed similar results, but with lower effect size or consistency for ChIP-seq data from different studies (Supp. Fig. 4c, Table S7), which could be attributed to the likelihood that some non-CCRs outside the gene-containing TADs may function as CCRs for other genes not tested in our experiments. Our findings demonstrate that CCRs are enriched in POU sequence motifs and associated with the enriched presence of OCT4, NANOG, QSER1 and TET1. Furthermore, CCRs, as defined by our functional assays, represent regulatory elements with distinct chromatin features that distinguish them from the previously defined primed enhancers or pre-enhancers.

### Pioneer TF FOXA2 preferentially binds to CCRs

Our results indicate hESCs have predefined regions that are competent to induce the transcription of lineage genes when interrogated with a targetable pioneer TF such as the iSAM system. However, it is unclear whether the chromatin features enriched at CCRs have a role in the activity of endogenous master TFs during lineage specification. We hypothesized that the chromatin features of CCRs may facilitate the binding of the master TFs during the onset of lineage specification. To test this hypothesis, we utilized CRISPRa to activate the expression of a well-known pioneer TF FOXA2, which plays critical roles in the development of endodermal organs (*25, 33, 34*). We expressed FOXA2 while keeping the cells in the E8 culture condition to mimic the earliest stages of lineage specification when a pioneer TF is expressed but before the cells acquire a new lineage identity. Through monitoring the expression of FOXA2 and DE differentiation markers daily, we identified 24 hours after doxycycline treatment as the optimal time window for analysis, with ∼60% of hESC expressing FOXA2 without activating DE TFs such as GATA6 (Fig. 5a). ChIP-seq revealed that in hESCs, FOXA2 mainly binds to regulatory genomic regions (introns and distal intergenic regions) and the binding sites are enriched in FOX motifs as expected (Supp. Fig. 5a-b). After filtering the FOX motifs, we found that FOXA2-bound regions were also enriched in the motifs associated to pluripotent factors SOX2, NANOG and OCT4 (Fig. 5b). To identify if the chromatin features enriched in the CCRs are associated to the regions where FOXA2 binds in hESCs, we generated meta-signal plots of the main factors that characterize CCRs, OCT4, NANOG, QSER1 and TET1 (ONQT) and checked their binding levels in FOXA2-bound regions (distal intergenic or intronic) that lack the H3K27ac mark, representing regions genomic loci with the potential to acquire enhancer functionality. We found significantly higher binding levels of the ONQT factors in FOXA2-bound regions compared to regions with FOXA motifs not bound by FOXA2 (Supp. Fig. 5c). A further analysis of the FOXA2 ChIP-seq signals in subsets of its bound regions, categorized based on the presence of any of the ONQT combinations (Fig. 5c), showed that the concomitant presence of the four ONQT factors was associated to significantly higher FOXA2 binding levels compared to regions bound by only one or none of the ONQT factors (Fig. 5d, Supp. Fig. 5d). This indicates that FOXA2 preferentially binds to regions with predictive features of CCRs based on the presence of ONQT. Considering the substantial overlap between CCRs identified in hESCs and enhancers in differentiated cells, we investigated whether the presence of ONQT correlated with the probability of FOXA2-bound regions in hESCs later becoming active enhancers in DE cells. We found that FOXA2-bound regions with the concomitant presence of all ONQT factors in hESCs had a significantly higher percentage of overlapping with FOXA2-bound active enhancers in DE cells (based on H3K27ac signal), compared to any of the other binding combinations (Fig. 5e). In summary, our data suggests that the combined presence of OCT4, NANOG, QSER1 and TET1 could facilitate further binding of master lineage TFs in the earliest stages of hESC lineage specification, thus creating a favorable environment for the subsequent establishment of active enhancers in differentiated cells.

**Figure 5.**
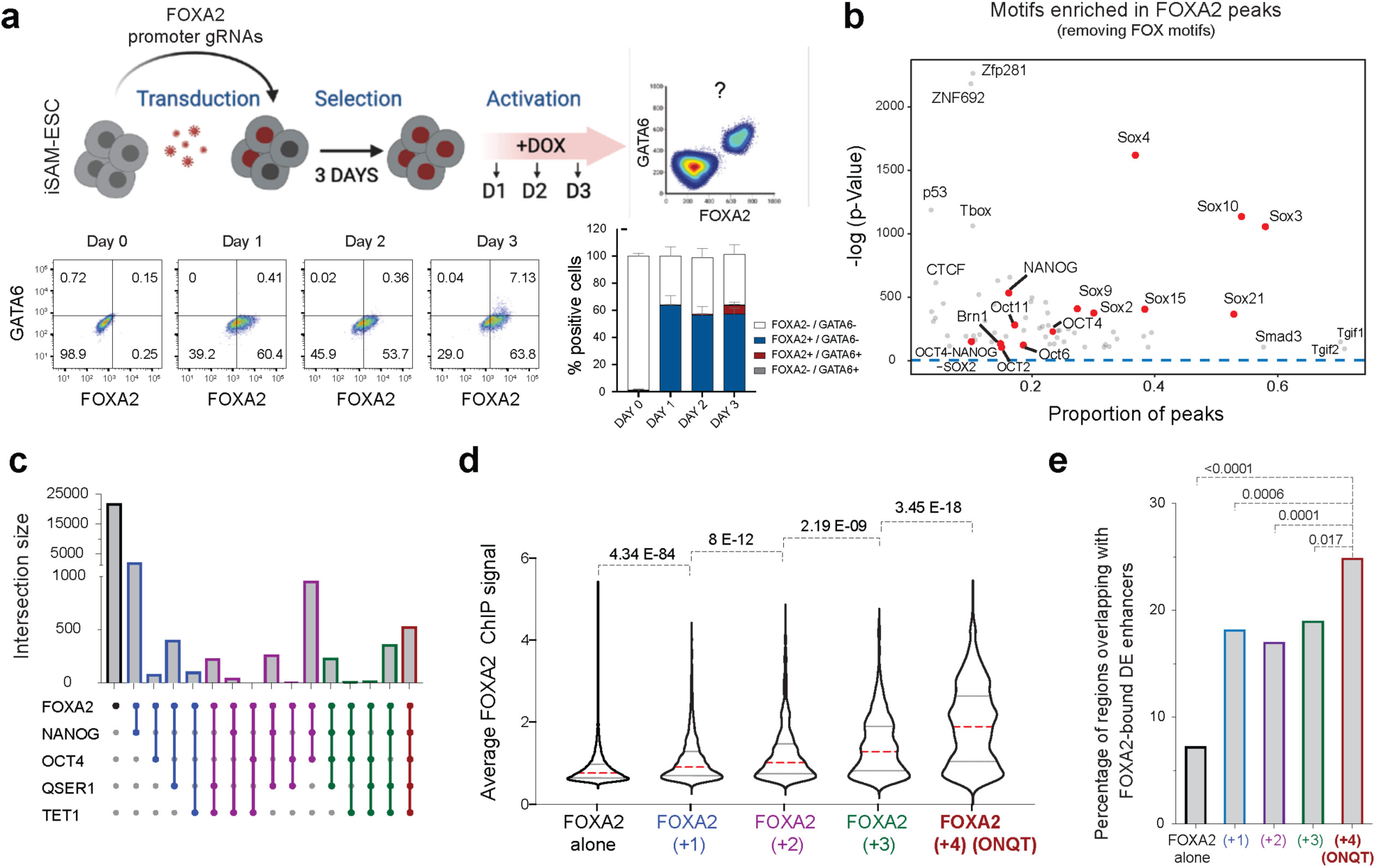
Pioneer TF FOXA2 binding profile in hESCs. **a)** Experimental design to determine the time-window to detect FOXA2 expression upon activation with the SAM tool targeting its promoter region at the ESC stage (Upper panel). Representative FACS dot plots and quantification of FOXA2 and GATA6 positive cells in ESCs upon FOXA2 activation with the SAM tool (lower panel). **b)** HOMER motif analysis plot to summarize the significantly enriched motifs (above the blue-dotted line) in the FOXA2 peaks detected upon expression at the ESC stage (after filtering out the FOX specific motifs). **c)** Upset plot to illustrate the intersection size between the FOXA2 peaks in hESCs and the main four factors enriched in CCRs (NANOG, OCT4, QSER1 and TET1). **d)** Mean ChIP FOXA2 score in the chromatin regions that overlap with any of the combination of the main four factors enriched in CCRs (Wilcoxon rank-sum adjusted p-values are shown). **e)** Percentage of FOXA2-bound regions in hESCs that overlap with active enhancers in DE cells (defined by ChIP-seq data) categorized based on their overlap with any of the combinations of the four factors enriched in CCRs (ONQT), Fisher’s exact test value are shown.

### Protection of CCRs from DNA hypermethylation by TET1 and QSER1 supports CCRs’ transcriptional competence

We identified that CCRs are regions able to induce the expression of lineage genes in hESCs, found within the TAD of their linked genes, and characterized by the enriched binding of OCT4, NANOG, QSER1 and TET1. To gain insight about the mechanism that supports the transcriptional competence of these regions we studied the role of QSER1 and TET1 in the CCRs. It has been previously shown that QSER1 and TET1 cooperate to keep DNA methylation valleys lowly methylated in hESCs (*32*). To test whether these factors had a similar effect in CCRs, we analyzed the DNA methylation levels at the CCRs and non-CCRs in previously generated hESC knockout (KO) lines for *TET1*, *QSER1*, and double KO (DKO) for *TET1*/*QSER1*, along with WT control cells (*32*) (Fig. 6a). The loss of both QSER1 and TET1 caused a significantly increase in the percentage of methylated CpGs at the CCRs but did not affect the methylation of the non-CCRs (Fig. 6b). To further dissect the functional consequences of having increased DNA methylation levels at the CCRs in *TET1*/*QSER1* DKO hESCs, we transduced the iSAM lentiviral vectors into the DKO hESC line and its matching WT control line. After confirming that both lines had comparable expression levels of the targetable activators used in the SAM system (Supp. Fig. 6a), we transduced the cells with representative gRNAs targeting several validated CCRs and after seven days we measured the expression of the associated genes by flow cytometry (Fig. 6c). We observed a significant decrease in the activation of the genes associated with the interrogated CCRs in the DKO cells compared to the WT control (Fig. 6d, Supp. Fig. 6b). Our findings demonstrate that QSER1 and TET1 promote the transcriptional competence of CCRs by protecting them from DNA hypermethylation.

**Figure 6.**
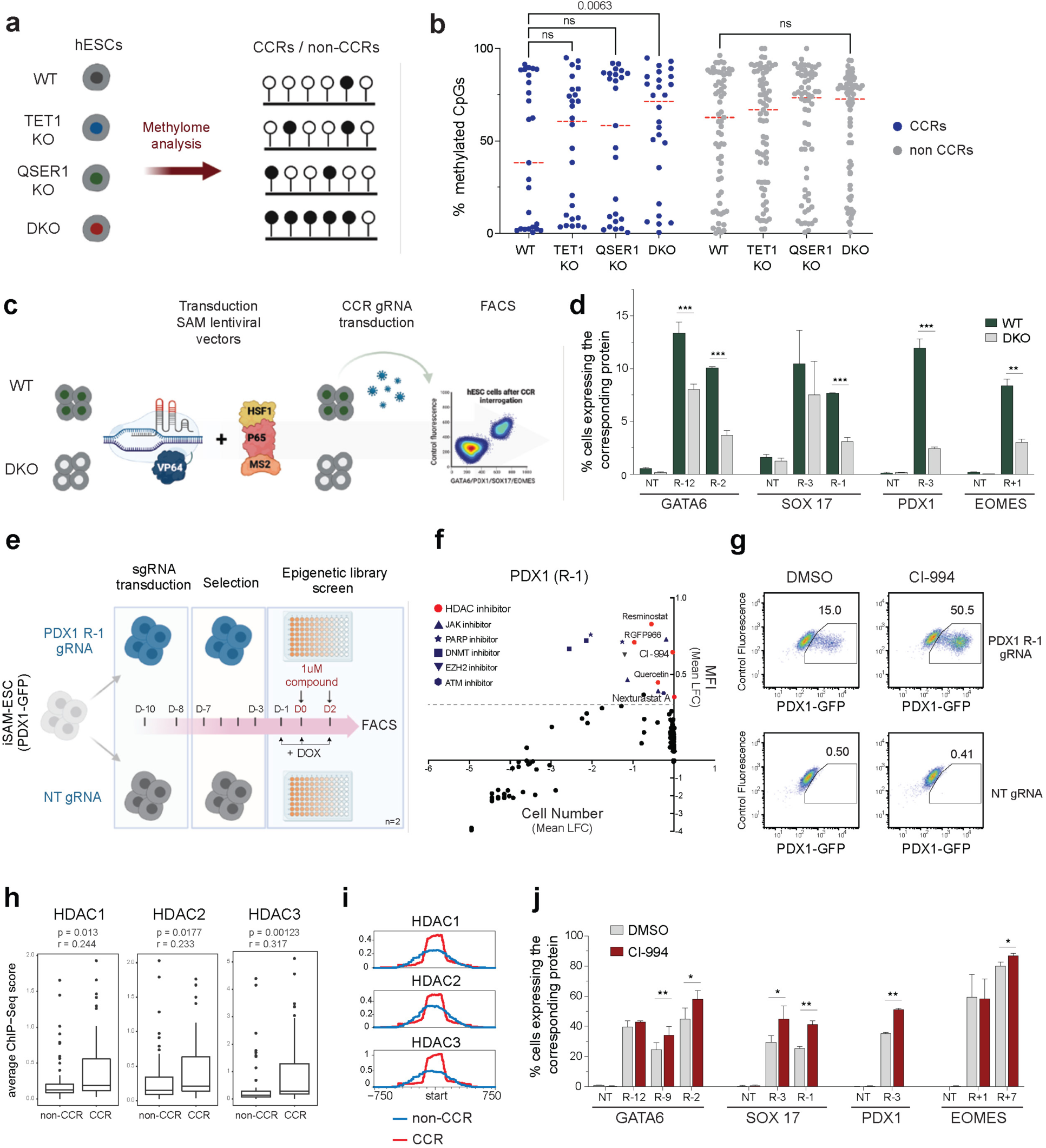
CCRs mechanism revealed by functional assays and epigenetic compounds screen. **a)** Experimental design to quantify changes in the DNA methylation levels around CCRs using the EPIC-methyl assays in hESC lines knocked out for TET1, QSER1 and double TET1/QSER1-KO (DKO) lines compared to WT control. **b**) Percentage of methylated CpGs at the CCR and non-CCR extended regions (see methods). Red dotted line indicates the median (FDR-adjusted p-value is shown). **c)** Experimental design to assess the transcriptional competence of several CCRs via CRISPRa interrogation, in hESC lines knocked out for QSER1 and TET1 compared to WT controls. **d)** Percentage of cells expressing the associated genes to each CCR after CRISPRa interrogation in WT and DKO cells measured by flow cytometry (based on protein expression). **e)** Experimental design followed to perform a chemical compound screen to identify epigenetic regulators of CCR’s transcriptional competence. **f)** Scatter plot that summarizes the results of the epigenetic screen. Each dot corresponds to the average LFC of the PDX1 mean fluorescence intensity (MFI) and the number of cells for each molecule of the library compared to DMSO (see Methods). Epigenetic modulators that significantly increase PDX1+ levels are indicated above the dotted line (n=2). **g**) FACS dot plots to illustrate the effect of the HDAC1 family inhibitor CI-994 after interrogation of the PDX1 CCR-1 with the SAM tool, compared to DMSO (left) and non-targeting gRNA controls (lower panel). **h**) Average binding of HDACs 1-2-3 measured by ChIP-seq score at the CCRs and non-CCRs in hESCs (Wilcoxon rank-sum test p values and effect sizes indicated on top). **i)** Average of normalized counts visualized as a MetaPeak using HDAC1-2 and 3 ChIP-signal data sets in the CCRs (red) compared to non-CCRs(blue). **j**) Percentage of cells expressing the associated genes to each CCR after CRISPRa interrogation of multiple CCRs and treatment with the HDACi CI-994 (red) vs DMSO (grey). Percentage of cells expressing the associated genes to each CCR interrogated is measured by flow cytometry of protein expression after molecule treatment and iSAM activation (n >3).

### HDAC1 family members are enriched at CCRs and regulate their transcriptional competence

The detection of chromatin features associated with CCRs is limited by the available ChIP-seq data. To further investigate chromatin-associated factors that could regulate CCRs, we performed an epigenetic screen employing a focused library containing ∼170 molecules that target various epigenetic pathways (Table S8, see methods). We transduced iSAM cells with either a gRNA targeting a representative CCR (*PDX1-*R-1) or a control NT gRNA. The transduced cells were then seeded into individual wells, where they were subjected to SAM activation as well as treatment with the library compounds or DMSO for three days (Fig. 6e). The NT control cells did not activate PDX1 expression, except when treated with a PARP inhibitor that strongly impairs cell growth (Supp. Fig. 6c). In contrast, the *PDX1*-R-1 gRNA transduced cells activate PDX1 expression as expected, and treatment with the epigenetic compounds caused both an increase and decrease in the percentage of PDX1+ cells compared to DMSO. To account for off-target effects due to a negative impact of the compound treatment on cell growth, we measured cell proliferation after treatment and found that all compounds that decrease PDX1 activation severely impair cell growth (Fig. 6f). In contrast, our screen revealed that treatment with multiple inhibitors of the histone deacetylases (HDACs) of the HDAC1 family significantly increase PDX1 activation without affecting cell growth (Fig. 6f). Among the tested compounds, the HDAC1-family inhibitor Tacedinaline (CI-994) demonstrated the strongest positive effect on PDX1 expression without compromising cell viability (Fig. 6g, Supp. Fig. 6d). To test whether this effect was associated to the enriched presence of HDAC1 family members at the CCRs, we performed ChIP-seq assays in hESCs to determine the binding profile of the histone deacetylases HDAC1, HDAC2 and HDAC3. Remarkably, we observed that all three HDAC1 family members were significantly enriched at CCRs when compared to non-CCRs (Fig. 6h-i). To test if HDAC1 family members regulate the transcriptional competence of CCRs beyond *PDX1-*R-1, we set up experiments to interrogate the activity of 8 CCRs of different genes (Supp. Fig. 6e). Strikingly, we observed that the CI-994 treatment caused a significant increase in the activation efficiency of CCRs for all the examined genes, as determined by flow cytometric analysis (Fig. 6j, Supp. Fig. 6f). Our findings show that CCRs are not only bound by chromatin activators such as QSER1 and TET1, but also by repressors represented by the HDAC1 family members. These factors exert opposing regulatory roles, and the delicate balance between activators and repressors help maintain CCRs in a poised state ready for rapid activation upon appropriate differentiation cues. While the molecular mechanisms may differ, the chromatin characteristics observed in CCRs share remarkable similarities with bivalent promoters, which are also regulated by opposing chromatin features to maintain a poised state for gene activation (*2*).

## Discussion

Our study has utilized a CRISPRa system to uncover a unique type of distal regulatory regions called CCRs. These CCRs exhibit a remarkable capability to activate the expression of lineage-specific genes in hESCs at levels comparable to those observed in differentiated cells. The intrinsic transcriptional competence exhibited by CCRs represents a distinct mechanism supporting the developmental plasticity required for ESCs to promptly activate lineage-specific genes at early stages of cell fate specification before the establishment of complete enhancer regulatory networks. Previous studies have alluded to a similar concept, proposing that certain chromatin regions in undifferentiated ESCs may exhibit distinctive features that prepare them for subsequent activation as enhancer elements in differentiated cells. For example, the histone variant H2A.Z and the low DNA methylation levels have been implicated in facilitating the binding of lineage TFs at the pre-enhancer regions in ESCs, while the H3K27me3 modification is believed to prevent premature activation of these regions (*4-6, 8, 10*). However, both H2A.Z and H3K27me3 modifications were equally enriched in CCRs and non-CCRs. We speculate that within the relatively broad genomic regions marked with histone modifications associated with pre-enhancers in ESCs, there exists a distinct class characterized as CCRs in our work. These CCRs can be distinguished by their exceptional capacity to activate gene transcription in the ESC stage.

Our study discovered several chromatin features that contribute to the transcriptional competence of CCRs. CCRs were found exclusively within the same TADs of their associated genes, but only a small fraction of them showed strong chromatin contacts with their target genes. This indicates that while some CCRs may exploit pre-established contacts to induce gene expression, most of them rely on indirect mechanisms of chromatin interaction, such as those proposed in the tracking/linking model (*28*). In addition, CCRs are enriched in antagonistic chromatin factors that can promote gene activation and repression. We established that the presence of the DNA demethylation factors TET1 and its partner QSER1 supports the transcriptional competence of CCRs by protecting them from excessive DNA methylation. Conversely, the presence of HDAC1 family members acts to prevent premature activation of CCRs. This “push and pull” mechanism resembles the bivalent domains found at the promoters of developmental genes and likely serves a similar purpose of preparing cells for the rapid activation of lineage-specific gene expression programs (*2*). In addition to bivalent-like chromatin features, CCRs also showed enriched binding of the pluripotent TFs OCT4 and NANOG. We found that the combined presence of OCT4, NANOG, QSER1 and TET1 premarks the regions that are preferentially bound by pioneer lineage TFs. Previous studies indicate pioneer TF binding is the initial event that triggers the formation of an enhancer element during the earliest stages of differentiation (*11, 33, 35*). Building upon our findings, we propose that the enriched binding of pluripotent TFs combined with DNA demethylation factors serves a “pre-pioneering” role by facilitating the binding of master lineage TFs. This concept is supported by a recent report suggesting that the chromatin regions with an increased presence of pluripotent TFs and TET1 in mESCs are enriched with future cell-type restricted enhancers (*36*). We predict that other factors such as FOXD3, which are enriched at regions where pluripotent TFs bind in ESCs and subsequently acquire enhancer-associated features in differentiated cells (*5, 37*), may also be associated with CCRs.

The ability of a genomic region to induce gene expression when interrogated with a CRISPRa system may depend not only on its intrinsic transcriptional competence but also on the strength of the transcriptional activators linked to the dCas9 molecule. We used the combination of the three activator domains in the iSAM system (VP64, HSF1 and p65) to discover CCRs in hESCs. In comparison, in a previous study on the *CD69* locus in T cells, the dCas9-VP64 system discovered CRISPRa-responsive elements only within regions already marked by H3K27ac (*15*). We speculate that employing stronger CRISPRa systems may enable the discovery of CCR-like regions also in T cells and other differentiated cells. This is supported by a recent report demonstrating the induction of β-like globin genes in HEK293 cells by targeting strong CRISPRa systems to a locus control region that functions as an enhancer in erythroid cells but lacks features associated to enhancers in HEK293 cells (*14*). Future work could determine whether antagonistic mechanisms, involving DNA demethylation factors and HDACs, also govern the regulation of CCRs in differentiated cells and adult tissues. Understanding these regulatory mechanisms beyond ESCs can provide insights into tissue homeostasis and regeneration processes. Moreover, the remarkable capability of CCRs to activate gene expression at levels comparable to differentiated cells presents exciting opportunities for directed reprogramming or trans-differentiation approaches aimed at generating specific cell types (*38*). On the flip side, exploring the involvement of CCRs in pathological conditions, such as cancer, could unveil how CCRs are hijacked to activate aberrant gene expression programs. Uncovering the roles of CCRs in such contexts may open avenues for therapeutic interventions, offering potential targets for precision medicine approaches.

## Supplementary Materials

- Methods

- Figures S1 to S6

- Tables S1 to S10 (additional files)

## Methods

### hESCs culture

Experiments in this study were performed using the hESC line H1 (NIHhESC-10-0043). hESCs were cultured in Essential 8 (E8) medium (Thermo Fisher Scientific, A1517001) on vitronectin (VTN) (Thermo Fisher Scientific, A14700) pre-coated plates at 37 °C with 5% CO2. Medium was changed every day. Cells were passaged every 4-5 days at a ratio 1:15-20 using 0.5 mM EDTA (KD Medical, RGE-3130) or TrypLE Select (Gibco, 12563029) to dissociate cells. 5 μM Rho-associated protein kinase (ROCK) inhibitor Y-27632 (Selleck Chemicals, S1049) was added into the E8 media when passaging or thawing hESCs. Cells were tested every 6 months by Memorial Sloan Kettering Cancer Center (MSKCC) Antibody & Bioresource Core Facility to verify they were mycoplasma-free. All experiments were approved by the Tri-SCI Embryonic Stem Cell Research Oversight Committee (ESCRO).

### CRISPRa and CRISPRi line generation

A H1 inducible Cas9 -PDX1-GFP line previously generated in our lab (*18*) which has a Cas9 cassette located at the AAVS1 locus was used to generate the CRISPRa and CRISPRi hESC lines. First, we aimed to substitute the endogenous Cas9 cassette by targeting the Cas9 protein to the Cas9 DNA flanking regions either with a dCas9-VP64 donor sequence from Addgene plasmid 61425 or a dCas9-KRAB donor. The dCas9-KRAB donor was generated through two rounds of QuikChange II Site-Directed Mutagenesis Kit (Agilent), using Puro-Cas9 donor (Addgene plasmid 58409) as template and the following primer pairs: RuvC1-F:R; RuvC2-F:R (Table S9). The final donor plasmid was generated through Gibson cloning (NEB) of a synthetic FseI-HA-KRAB-AscI DNA fragment (GenScript) flanked by 50 nucleotide homology arms into Puro-dCas9 donor cut FseI-AscI and the plasmid backbone was further replaced with one containing a low copy number origin of replication. PCR-amplified donor sequences were independently cloned by In-fusion reaction (Takara) into the backbone of the Addgene plasmid 58409 which contains the homology arms (HA) to target the AAVS1 locus. The resulting plasmids contained the HA sequences flanking the donor sequences, to be recombined with the endogenous AAVS1 sequence following a previously reported strategy (*39*) (Supp. Fig. 1a). In brief, iCAS9 -PDX1-GFP cells were seeded in VTN-coated plates using regular E8 medium and treated with doxycycline (Dox) 2 μg/ml for 2 days to activate Cas9 expression and for each final cell line 3x10^5^ cells were transfected with 1ug of the corresponding donor vector and 80ng of the targeting gRNAS (CR2R-CR2L gRNAs for the idCas9-VP64 line and CR1R-CR2L gRNAs for the idCas9-KRAB) (Table S9) using the Lipofectamine Stem Transfection Reagent (Thermo Fisher Scientific) following manufacturer’s guidelines. Transfected cells were treated an additional day with Dox, and treated 4 days with G418 (500 μg/ml) to select the clones with the RTTA cassette), recovered one day in basal E8 and treated with Blasticidin (10μg/ml) for another 4 days (to select the clones with the dCas9-VP64 or dCas9-KRAB cassette). Surviving colonies were dissected into single cells using TrypLE Select Enzyme (1X) and 1000 cells were seeded into individual VTN-coated 10cm plates. Individual colonies were manually picked, grown for one week and tested for genotyping using the sequencing primers listed in table (Table S9). Clones with the correct sequences were further validated by treatment with Doxycycline over 3 days and analysis of the expression of the dCas9-KRAB or dCas9-VP64 cassettes by qPCR, Western blot and FACS. After several rounds of verifications, we obtained one positive clone for the dCas9-KRAB line and two clones for the dCas9-VP64 line. We picked one clone for the dCas9-VP64 line together with the dCas9-KRAB and both clones were karyotyped to verify absence of genetic abnormalities by the MSKCC Molecular Cytogenetics Core. To generate the iSAM line, we transduced the idCas9-VP64-PDX1-GFP hESC line with the Addgene plasmid 61426, containing the EF1-MS2-P65-HSF1-hygro cassette. After 4 days of selection with hygromycin (200 μg/ml), 1000 cells were seeded in 10cm plates and individual clones were picked and grown separately. Three rounds of clonal selection were performed, and 12 final clones were treated with doxycycline for 3 days and tested for MS2 and dCas9 protein expression by flow cytometry. The clone with the highest expression of both was chosen as the iSAM hESC line.

### gRNA libraries

For the PDX1 CRISPRa and CRISPRi screens, we designed a tiled library using the GuideScan package (*40*) covering 50kb upstream from TSS (2998 gRNAs), PDX1 intronic regions (698 gRNAs) and 50kb downstream from 5’ end of the 2^nd^ exon (3442 gRNAs), as positive controls we included gRNAs targeting exonic regions (99 gRNAs), and as negative controls non-targeting to human genome (200 gRNAs) and safe-harbours for each chromosome (200 gRNAs). The gRNA library targeting the candidate enhancers surrounding *GATA6*, *EOMES*, *SOX17* and *MIXL1* was derived from a previously published library(*26*). In brief, regions 2mb upstream/downstream around the 4 TFs that showed any chromatin accessibility at the ESC or DE stage were selected for interrogation, removing promoters and exons. 163 regions were selected to design fully tiled gRNAs using CHOPCHOP(*41*). In addition, 1100 gRNAs targeting safe harbor loci and 3 gRNAs targeting the *SOX17* promoter were included as negative and positive controls respectively. gRNA oligos were synthesized on-Chip (Agilent). PDX1 synthesized oligos were amplified and restriction cloned into lentiGuide-puro (Addgene; 52963) and lenti MS2 grna-Puro (Addgene; 52963), while the second library was only cloned into the lenti MS2 grna-Puro backbone by the MSKCC Gene Editing & Screening Core Facility. Cloned plasmid libraries were PCR amplified to incorporate adapters for NGS. Samples were purified and sequenced using Illumina HiSeq 2500 platform. FASTQ files were clipped by position and reads were mapped back to the reference library file to show relative abundance of reads per gRNA. Reads within each sample were normalized to total number of mapped reads and library size. The Overall Representation of the libraries was charted over a one-log fold change to evaluate if any gRNA was over- or under-represented in the final library.

### Lentiviral gRNA libraries

gRNA libraries lentiviral particles generation was performed as previously described (*42*). In brief, a total of 9.45 μg of each library plasmids combined with 6.75 μg lentiviral packaging vector psPAX2 and 1.36 μg vesicular stomatitis virus G (VSV-G) envelope expressing plasmid pMD2.G (Addgene plasmids 12260 and 12259) were transfected with the JetPRIME (VMR; 89137972) reagent into 1x10^6^ 293T cells in a 10cm plate to produce the lentiviral particles. Fresh medium was changed 24h after transfection and viral supernatant was collected, spun at 1000 rpm for 5 minutes, filtered, and stored at −80°C, 72h after transfection.

### CRISPRa screen for PDX1 CCRs

The PDX1 lentiviral library with the MS2-gRNA backbone was transduced into 12 million hESC iSAM cells distributed in 10 x 10 cm plates, while the library cloned using the regular lenti-gRNA backbone was transduced into 12 million hESC dCas9-VP64 cells distributed in 10 x 10 cm plates, to reach a MOI =0.3. Cells were reverse transduced in the presence of ROCK inhibitor (RI) (10 μM, Selleck Chemicals) and protamine sulfate (PS) (6 μg/ml, MP Biomedical). Both transduced cell lines were kept under puromycin selection (1 μg/ml) for 4 days, and after one day of recovery with regular E8 media, were grown with Dox during four additional days. The presence of cells expressing PDX1 (reported by GFP) was verified by flow cytometry, after being stained with LIVE/DEAD reagent (Invitrogen, Catalog# 34955) for 15 mins at RT. All the iSAM-hESCs expressing PDX1 (GFP+) cells were FACS-sorted using FACSaria sorters (BD Biosciences) by the MSKCC cytometry facility, a corresponding number of PDX1-GFP (-) cells were also sorted, pelleted down and kept at -80 for downstream gRNA sequencing. The screen was performed twice using independent biological replicates.

### CRISPRi screen for PDX1 enhancers

The PDX1 lentiviral library with the regular lenti-gRNA backbone was reverse transduced into 12 million hESC (i)dCas9-KRAB hESCs distributed in 10 x 10 cm plates following the previous conditions (MOI =0.3, PS =6 μg/ml, RI =10 μM). Cells were kept under puromycin selection for 4 days, and after one day of recovery with regular E8 media. 12 million selected cells were seeded into 6well plates (4x10^5^ per well) in VTN-coated plates, grown for 48 hours in E8 medium with Dox. After washing with Phosphate Buffered Saline (PBS) w/o Ca2+ & Mg2+, cells were differentiated into Pancreatic Progenitor (PP) cells utilizing a protocol described previously(*22*) (see *hESC differentiation* below). Doxycycline treatment was administered throughout the nine-days differentiation process. After, cells were stained with LIVE/DEAD reagent and FACS-sorted using FACSaria sorters (BD Biosciences) based on PDX1 expression (reported by GFP expression) by the MSKCC cytometry facility. PDX1 positive and PDX1 negative cells were sorted aiming for a minimum representation = 500x (number of cells with a unique sgRNA). Cells were pelleted down and kept at -80 for downstream gRNA sequencing. The screen was performed twice using independent biological replicates.

### Expanded CRISPRa screen

The lentiviral library targeting the EOMES, GATA6, SOX17 and *MIXL1* loci cloned into the MS2-gRNA backbone was transduced into 12 million hESC iSAM cells distributed in 10 x 10 cm plates, to reach a MOI =0.3. Cells were reverse transduced in the presence of RI (10 μM) and PS (6 μg/ml). After one day of recovery, the transduced cells were kept under puromycin selection and doxycycline for 4 days in E8 regular media, and then 2 additional days with E8 media and Dox. After, ∼ 60 x10^6^ cells were divided into two equal batches, stained with LIVE/DEAD reagent, washed using FACS Buffer (PBS, 2.5% Fetal bovine serum and 0.5mM EDTA), pelleted down at 500 RCF for 5 mins and fixed using eBioscience™ Fixation/Permeabilization Concentrate (Invitrogen) and Diluent Solution (Invitrogen) at RT for 1 hour. Following fixation, one batch was with incubated with GATA6-PE and SOX17-FITC and the second one with EOMES-FITC and MIXL1combined with mouse-647 secondary antibody (Table S10) at RT in Fix&Stain 1X Permeabilization buffer solution (eBioscience). As a control to create the gating strategy for sorting, 1x106 cells were transduced with a NT-gRNA and kept under the same conditions as above. All the positive cells for the GATA6+/SOX17+ and the EOMES+/MIXL1+ fractions were sorted and a corresponding number of GATA6-/SOX17- and EOMES-/MIXL1-were also sorted by the MSKCC cytometry facility, the positive and the negative fractions were individually combined, pelleted down and kept at -80 for downstream gRNA sequencing. The screen was performed twice using independent biological replicates.

### gRNA sequencing

The gRNA enrichment sequencing was performed by MSKCC Gene Editing & Screening Core Facility as previously described (*18, 26*). Briefly, genomic DNA from sorted cell pellets was extracted using the QIAGEN Blood & Cell Culture DNA Maxi Kit (QIAGEN; 13362) and quantified by Qubit (Thermo-Scientific) following the manufacturer’s guidelines. For the CRISPRa expanded screen, cells were previously decrosslinked and treated with proteinase K (20mg/ml). A quantity of gDNA covering the representation of gRNAs described above was amplified with oligos containing the Illumina adapters and multiplexing barcodes by PCR. Amplicons were quantified by Qubit and Bioanalyzer (Agilent) and sequenced on the Illumina HiSeq 2500 platform. The gRNA library sequences were used to align the sequencing reads and the counts for each gRNA were determined.

### Preliminary analysis to validate CRISPRa screen targeting PDX1 locus

A preliminary analysis for the CRISPRa screen to detect *PDX1* CCR’s was performed to ensure the positive gRNA controls worked. We selected the gRNAs overlapping the promoter region of *PDX1* (400bp upstream-200 bp downstream from the TSS) and calculated the differential gRNA read abundance between the PDX1+ and the PDX1-cell populations (measured as Log Fold Change). The average LFC of the positive gRNA controls was used as a threshold to determine the number of gRNAs enriched in the PDX1+ fraction (Table S1).

### Sliding window analysis for CRISPRa and CRISPRi

#### Read count normalization and computing fold changes

To render the counts from different replicates comparable, the values are adjusted using the median ratio method (*20*). Let *c*_g_^+^and *c*_g_^-^ denote the read counts of gRNA g for each positive and negative sorted fraction in a replicate, respectively. The size factor is calculated as the median of size factors for individual gRNAs:

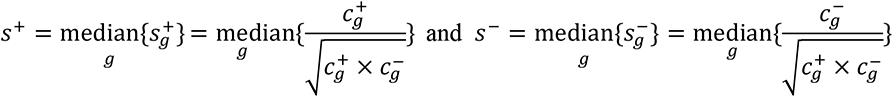

Then, for gRNA *g*, the log fold change between the positive and the negative sorted fraction is derived from the adjusted counts:

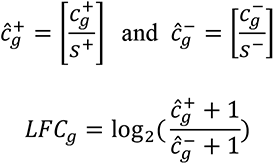

#### Calculating the moving average

The moving average of log fold change values is calculated using the sliding window approach (*21*). After ordering the gRNAs by their genomic coordinates, the average LFC is computed over a window of *N* gRNAs, sliding by one at a time. Given replicates I and II, let 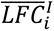 and 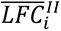 denote the average *LFC* of the *i*th window in each replicate. With *N* ranging between 1 to 50, the window size resulting in the highest Pearson correlation coefficient between *L̄F̄C̄*^I^ and *L̄F̄C̄*^II^ is selected. For visualization purposes, the *LFC* of a screen across the genome is represented by the mean between replicates as 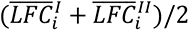.

#### Quantifying significance and reproducibility

The significance of window *i* in each replicate is quantified with the *Z*-score of its *LFC* as *Z_i_* = 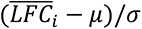 where the mean and standard deviation, respectively *μ* and *σ*, are calculated from *LFC* of individual gRNAs in that replicate. Reproducibility between the replicates is measured by the reverse of Irreproducible Discovery Rate (*IDR*)(*43*). Using *R* function *idr::est.IDR*, the rate for window *i* can be estimated by *IDR_i_* between |*Z_i_*^I^| and |*Z_i_*^II^|, which represent its significance in replicates I and II.

### Identifying candidate regulatory regions for PDX1

The following analysis was performed on CRISPRa and CRISPRi screens, respectively, to discover the candidate PDX1 regulatory regions. First, any gRNA with < *min_count* (=50 for CRISPRa and 20 for CRISPRi) reads in either replicate was excluded. For each replicate, log fold change values between GFP^+^ and GFP^-^ were calculated as described in *“Read count normalization and computing fold changes”*. To normalize read counts in this analysis, the size factors were calculated as the median of size factors for non-targeting gRNA controls (i.e., 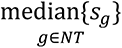). To mitigate the noise of log fold change values, the moving average was computed for a sliding window of *N* (= 30 for CRISPRa and 10 for CRISPRi) gRNAs (see *‘Calculating the moving average’*). Then, the significance and reproducibility of each window was quantified as described in *‘Quantifying significance and reproducibility’*. To calculate the Z-scores in this analysis, the mean and standard deviation were derived from log fold change values of non-targeting gRNA controls (i.e., *LFC_g_* for *g*∈*NT*). Significant windows were selected based on their length (*i.e.*, span from the starting coordinate of the window’s first gRNA to the ending coordinate of its last), significance in each replicate, and reproducibility between the replicates: window *i* was discarded if length*_i_*>*max_length* or |*Z_i_*^I^| <*min_score*or |*Z_i_*^II^| <*min_score* or *IDR_i_*>*max_IDR* (*max_length*=250bp, *min_score*=0.25, *max_IDR*=0.05 for CRISPRa and *max_length*=200bp, *min_score*=3, *max_IDR*=0.001 for CRISPRi). Finally, we created the candidate regions by merging the genomic regions corresponding to the selected windows in an iterative process, as follows: out of all consecutive regions whose combination would span <*max_span* (= 1000bp for CRISPRa and 2000bp for CRISPRi), the pair with the minimum distance is combined; if no such pair exists, stop and report the final set of genomic regions. The coordinates of the identified regions are summarized in Table S2. For comparative analyses purposes, the PDX1 non-CCRs were randomly chosen from the regions that did not meet the criteria for CRISPRa (*i.e.*, window *i* for which |*Z_i_*^I^| <0.25 or |*Z_i_*^II^| <0.25 or *IDR_i_*>0.05) and had an average length equal to the PDX1 CCRs, see Table S3.

### Expanded CRISPRa screen analysis to identify CCRs for GATA6, EOMES, SOX17 and MIXL1 at the ESC stage

To identify the CCRs associated to *GATA6, EOMES, SOX17* and *MIXL1*, the gRNAs sequencing reads for the positive (*i.e.*, cells expressing any of the 4 genes after CRISPRa interrogation) and the negative FACS sorted fraction were obtained by NGS, and the following analysis was performed for each gene respectively. First, off-target gRNAs were filtered out using CRISPOR (*44*), as previously reported (*26*). For each replicate, after adding *pseudocount*=1000 to the read counts, log fold change values between the positive and the negative sorted fractions were calculated as described above (“*Read count normalization and computing fold changes”*). For this analysis, the log fold change values of individual gRNAs were used instead of the moving average; in other words, window *i* is the *i* th gRNA and the notation in *‘Calculating the moving average’* holds for *N*=1. Then, the significance and reproducibility of each gRNA was quantified as described in *‘Quantifying significance and reproducibility’*. Significant gRNAs were selected based on their significance in each replicate and reproducibility between the replicates: gRNA *i* is discarded if *Z_i_*^I^<1 or *Z_i_*^II^<1 or *IDR_i_*>0.05. We defined a genomic region for each group of nearby significant gRNAs, extended until there were none closer than 100bp. Finally, we created the CCRs by extending the resulting regions by 150bp around the center and merging the overlapping regions. We identified non-CCRs by subtracting the CCRs from the differential ATAC peaks between the ESC and DE stages that were targeted by the screen: ATAC peaks and parts of the ATAC peaks that did not overlap CCRs and were longer than 150bp were considered non-CCRs. gRNA enrichment for each interrogated region and the final list of CCRs and non-CCRs are provided in Table S3.

### Validation for individual CCRs by flow cytometry

CCRs and non-CCRs (for the extended CRISPRa screen) were determined based on the previous methods and individual representative gRNAs targeting those regions were chosen for individual validations. In brief, the selected gRNAs (Table S9) for each candidate CCR were cloned into the lentiguide-Puro for validation of the CRISPRi screen and the lentiguideMS2-puro for the CRISPRa screens. For lentivirus packaging, 0.65 μg of lentiGuide-puro (or lentiguideMS2-puro), 0.1 μg of pMD2.G, and 0.4 μg of psPAX2 plasmids were transfected into 5x10^5^ 293T cells using JetPRIME (89137972; VMR) reagent. Viral supernatants were harvested as previously described. For experiments to validate CCRs using CRISPRa we individually transduced 2x105 iSAM cells with gRNA lentiviruses to reach a MOI = 0.3 in a well of a 12wp. Assays included the transduction with and additional non-targeting gRNA as negative control (Table S9), after one day of recovery we kept the cells under puromycin (1μg/ml) treatment in regular E8 media and added doxycycline for 72-96 hours, changing the media every day. Cells were rinsed with PBS w/o Ca2+ & Mg2+ and treated with TrypLE Select Enzyme (Invitrogen, Catalog# 12563-029) for 2-3 mins for dissociation and collected using FACS Buffer. The cell suspension was centrifugated at 500 RCF for 5 mins, fixed and stained with LIVE/DEAD and specific antibodies as described in *Expanded CRISPRa screen*. For validation of gRNAs from the CRISPRi screen, we performed an equivalent assay. We transduced 2x10^5^ idCas9-KRAB cells with individual gRNAs lentiviruses and kept the cells under puromycin selection for 3 days (after one recovery day). Cells were collected and counted to differentiate them to the pancreatic progenitor stage, following the protocol described in *hESC differentiation to DE and PP*. Cells were kept in doxycycline treatment during the 9 days of differentiation. At day 9 cells were collected, LIVE/DEAD stained and analyzed the PDX1-GFP expression by flow cytometry. To further validate GFP signal as a reporter of the PDX1 expression, control experiments were performed, fixing cells after differentiation, and staining with a PDX1 antibody (Table S10).

### Validation for individual CCRs by Immunofluorescence staining

iSAM or idCas9-KRAB hESCs were transduced with individual gRNAs targeting candidate CCRs and non-targeting controls as described above. Adherent cells (at ESC stage for the CRISPRa validations and PP stage for the CRISPRi validations) were rinsed once with PBS with Ca2+ and Mg2+ (PBS++) and fixed with 4% Paraformaldehyde (Electron Microscopy Sciences 32% Paraformaldehyde, #15714-S, diluted in PBS++) for 10 mins. Following fixation, cells were rinsed once with PBS++ and washed 3 times with PBST (0.1% Triton X (Sigma Aldrich, # T9284) in PBS w/o Ca2+&Mg2+) in a rocking shaker for 5 mins. After incubation with a with blocking solution (Donkey Serum) for 5 mins at RT, cells were incubated with primary antibody diluted in blocking solution overnight at 4C with gentle rocking. Following ON incubation, cells were rinsed once in PBST, washed 3 times with PBST in a rocking shaker for 5 mins, and incubated with a secondary antibody for 1 hour at RT with gentle shaking. After an intermediate rinse step with PBST, the cells were stained with DAPI (conc) for 15 mins at RT with gentle shaking, washed 3 times with PBST in a rocking shaker for 5 mins and stored at 4C in dark until imaging with PBS++ solution. Images were generated using the Confocal Laser Scanning Platforms Leica TCS SP5 and SP8. Acquisition parameters were fixed across experiments that involved comparison of the protein expression levels among multiple samples. Max. projections of Z-stacked individual images were generated with Fiji (ImageJ) and Imaris 9.6.

### Clonal enhancer KO hESC line generation for CCR validations

Enhancer KO hESC lines were generated with paired crRNAs that flanked *PDX1* region -7 and region -1 (Table S9). crRNAs and tracrRNA were ordered as RNA oligos from IDT. hESCs were treated with 2-μg/ml doxycycline and 10-μM ROCK inhibitor Y-27632 for 24h. Cells were then dissociated with 1X TrypLE Select and transfected for a final concentration of 0.01-μM of each crRNA and 0.01-μM of tracrRNA with Lipofectamine Stem Transfection Reagent (Thermo Scientific) following manufacturer’s guidelines. Transfected cells were then cultured in E8 with 2-μg/ml doxycycline for 24h, and then in E8 for 24h. ∼2000 cells were seeded into a 10cm plate and after 5-8 days of culture single colonies were picked and expanded. Individual colonies were genotyped using the primers listed in Table S9, after genomic DNA extraction with the DNeasy Blood & Tissue DNA Kit (QIAGEN).

### hESC differentiation to DE and PP

Following a reported protocol (*22*), hESCs were seeded into VTN-coated 6well plates (4x10^5^ per well), grown for 48 hours in E8 medium. After washing E8 away with PBS, cells were cultured in S1/2 medium, supplemented with Activin A (100ng/ml) and CHIR99021 (Stemgent, 04-0004-10) (5 µM on the 1st day and 0.5 µM on the 2nd day) for three days. Cells reached the Definitive Endoderm (DE) at day 3 and were used for validation experiments (expression analysis of DE-related genes). To continue with the protocol to generate pancreatic progenitors, cells were washed with PBS and further cultured for two days in S1/2 medium supplemented with FGF7 (50ng/ml) (PeproTech, 100-19) and vitamin C (0.25mM) (Sigma-Aldrich, A4544). To promote final differentiation towards the PP stage, cells were cultured for two days in S3/4 medium supplemented with FGF7 (50 ng/ml), vitamin C (0.25mM), retinoic acid (1µM), LDN (100 nM) (Stemgent, 04-0019), SANT-1 (0.25 µM) (Sigma, S4572), TPB (200 nM) (EMD Millipore, 565740), and ITS-X (1:200) (Gibco, 51500056).

### Flow Cytometry analysis

Flow cytometry analysis was carried out according to a previously reported protocol (21). Antibodies utilized for flow cytometry are described in Table S10. Protein expression by flow cytometry analyses was performed using a BD LSR Fortessa or BD LSRII from the MSKCC core facility. Analysis of flow cytometry data was done using FlowJo v10.

### 4C

For each primary digestion reaction, 10 million hESCs were transferred into 15 ml Falcon tubes and fixed with 10 ml of 1% formaldehyde (Thermo Scientific, 28908) in PBS without Ca+/Mg+ for 10 min at room temperature (RT) (on a rotation device). After quenching with freshly prepared ice-cold 1M glycine, tubes were centrifuged for 5 min 500g at 4°C. Cells were kept on ice, washed with PBS, centrifuged for 5 min 500g at 4°C, and pellets were frozen in liquid nitrogen and stored at −80°C. Next, cells were vigorously resuspended in 1 ml of ice-cold lysis buffer (10 mM tris (pH 8), 10 mM NaCl, 0.2% NP-40, and 1 tablet of complete protease inhibitor (Roche)], transferred to 9 ml of prechilled lysis buffer, and incubated for 15 min on ice. Following centrifugation at 500g for 5 min at 4°C, the pellet was resuspended in 500uL 1.2x CutSmart buffer containing 0.5% SDS and incubated for 1 hr at 37°C while shaking at 750 rpm. After, SDS quenching was performed by adding 75uL 20% Triton X-100 and incubating for 1 hr at 37°C while shaking at 750 rpm. 10 uL of the sample were taken as an undigested control. The remaining samples were treated with DpnII enzyme (NEB, R0543M) (30ul) in CutSmart buffer (NEB, B7204S) (25ul) and samples were incubated ON at 37°C under agitation (750rpm). After first digestion, 5μl of the sample were taken as a digested control and the efficiency of the chromatin digestion was verified by electrophoresis, detecting a smear between 0.2 and 2 kb in a 1.5% agarose gel. DpnII was deactivated by (6565°C, 20 minutes, 750 rpm), and ligation of DNA ends between the cross-linked DNA fragments was performed in T4 ligation buffer (NEB, B0202), ATP (NEB, P0756S), 10% Triton X-100, 20mg/ml BSA and T4 DNA Ligase (NEB, M0202) ON at 16°C (gently rocking) followed by 30min at RT. Then, ligated samples were treated with proteinase K and reverse crosslinked overnight at 65C. Following RNase (Sigma-Aldrich, 10109142001) treatment, phenol/chloroform extraction and DNA precipitation, the pellets were dissolved in 10mM Tris pH 8. Then the efficiency of the DNA extraction was verified on a 1.5% agarose gel. For the second digestion, 5uL of NlaIII enzyme diluted in CutSmart buffer were added to the DpnII-ligated 3C template and samples were incubated ON (37°C, 750rpm). NlaIII was inactivated at 65°C for 20 min, and a second ligation was performed by adding T4 ligation buffer, 10mM ATP, T4 DNA Ligase, and ddH2O, incubating ON at 16°C. Ligation product was extracted by phenol/chloroform. DNA concentration of each digested sample was calculated using the Qubit brDNA HS assay kit (Invitrogen). For library preparation, viewpoint primers were designed either around promoter of EOMES, GATA6, SOX17 and PDX1, depending on the availability of the nearest restriction enzyme sites (see Table S5). Library preparation was then performed using 200 ng of 4C-template DNA with the Expand Long Range PCR kit (Millipore, 4829042001) under specific PCR conditions (94 °C for 2 min, 16 cycles: 94 °C for 10 s; 59°C for 1 min; 68 °C for 3 min, final extension: 68 °C for 5 min). Amplified DNA was pooled, and primers were removed using AMPure beads (Beckman Coulter, A63881). A second round of PCR was performed using the amplified DNA as a template and overlapping primers to add the P5/P7 sequencing primers and indexes (94 °C for 2 min, 20 cycles: 94 °C for 10 s; 60 °C for 1 min; 68 °C for 3 min and 68 °C for 5 min). The samples quantity and purity were determined using a Qubit fluorometer. 4C PCR library efficiencies were confirmed by Agilent Bioanalyzer and libraries were sequenced on a HiSeq4000 in PE150 mode by the MSKCC Genomics Core Facility. Three independent assays per viewpoint were performed.

### Hi-C Processing

Hi-C raw data from undifferentiated hESCs (https://data.4dnucleome.org, accession numbers: 4DNESDO2ZYBM, 4DNESQMUTYXH, 4DNESFL8KDMT)(*26, 45, 46*) was processed using HiC-Pro 2.11.4 (*47*). Hi-C-Pro utilized bowtie2 2.4.1 for alignment steps, samtools 1.8 for sorting, and Python 3.8.3 for scripting (*48, 49*). Reads were aligned to the analysis set of hg19, which excludes alternative contigs from the genome with default settings. After alignment, MAPQ30 reads were used for further analysis. Reads from technical repeats were combined and dededuplicated, while reads from biological replicates were simply merged. Reads for replicated and merged maps were converted into .hic files with Juicer 1.22.01 (*50*).

### TADs

Topologically associating domains were identified using TopDom 0.10.1 via the interface available in Hi-C-DC+ (*29, 51*). Prior to running TopDom, HiTC was used to carry out ICE balancing via the wrapper function provided in HiC-DC+ (*52*). ICE balancing was carried until convergence or 50 iterations. TAD calling was performed using 50 kb bins, scanning either 5 or 10 bins up and downstream to compute boundaries (see Table S4).

### 4C Processing

4C processing was first carried out with pipe4C v1.0 (*53*). Default settings were used for all analysis. After all reads were processed, the data was first re-binned into fixed bins of 2500 bp, and then a sliding window of 5 was used to aggregate counts. These smoothed counts were then used for subsequent analysis. A local model analogous to HiC-DC+ was used to identify local loop enrichments at an FDR of 0.1 (*29*). In brief, HiC-DC+ assumes a Negative Binomial GLM parametrized with a fixed design spline for the distance term. To deal with the intrinsic variation of 4C data, we adopted a Generalized Additive Model for Location Shape and Scale(*54*) that uses smooth spline terms for the distance dependence, and simultaneously models the dispersion parameter of the observations. These models were fit to 4C capture experiment to model local dependencies.

### 4C Scoring

To score events observed within each replicate, we used a z-scoring procedure based on quantile residuals derived from the local model analogous to HiC-DC+. After scoring, these quantile residuals should follow an approximately normal distribution, and so, 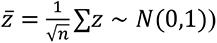 as well. These were then used to identify differences between the CCRs and non-CCRs. A kernel density estimator from the CRAN package sm 2.2-5.7.1 was used to estimate the KDE and its associated confidence band of equality (*55*).

### ChIP-seq assays

ChIP-seq was performed as previously described (*42*). Briefly, for each sample, around 30 million cells were crosslinked with 1% formaldehyde and quenched with 0.125M glycine. Fixed cells were collected in cold PBS buffer and washed with the same. Then the cells were snap frozen and saved at -80 °C. On the day of immunoprecipitation, cell pellet was thawed on ice, resuspended in 700uL SDS buffer (1% SDS, 10 mM EDTA, 50 mM Tris–HCl, pH 8), and incubated for 10min on ice. Sonication was performed on a Branson Sonifier 150 set at 30% amplitude for 5min30s (10s on/off pulsing). Supernatant was pre-cleared with Dynabeads and then incubated overnight with antibodies (10ug for transcription factors and HDACs, 5ug for H3K27ac, see Table S10) at 4°C.). On the next day, the ChIP samples were incubated with Dynabeads for 6 hours at 4°C. Then the beads were pelleted and washed twice with low salt (0.1% SDS, 1% Triton X-100, 2 mM EDTA, 20 mM Tris–HCl, pH 8, 150 mM NaCl), high salt (0.1% SDS, 1% Triton X-100, 2 mM EDTA, 20 mM Tris–HCl, pH 8, 500 mM NaCl), and TE buffer (10 mM Tris–HCl, pH 8, 1 mM EDTA) respectively. The DNA was eluted from the beads by incubating in elution buffer (1% SDS, 0.1 M NaHCO3) at 65°C for 15min and decrosslinked with 5M NaCl at 65°C overnight. A total of 10 μl 0.5 M EDTA, 20 μl 1 M Tris–HCl, pH 6.5, and 1 μl Proteinase K (20 mg ml−1) were added to decrosslinked product and incubated for 1 h at 45 °C. DNA was isolated by using QIAquick PCR purification kit (Qiagen, 28104; QIAGEN). Then the sequencing library was generated by the NEBNext® Ultra II DNA Library Prep Kit (New England Biolabs; E7103S) and NEBNext® Multiplex Oligos for Illumina® (New England Biolabs Index Primers Set 1; NEB, E7335S). Samples were pooled and submitted to MSKCC Integrated Genomics Operation core for quality control and sequencing on Illumina HiSeq 4000 platform.

### ChIP-seq analyses

Paired-end reads were mapped to hg19 reference genome with bowtie2 (*48*) using default parameters. Peaks were called using macs2 (Model-based Analysis of ChIP-Seq (MACS) (*56*), with the options to filter duplicates using –keepdup auto with the permissive p-value threshold of 0.01. Peaks were further filtered down using irreproducible discovery ratio (IDR) (*43*) filter of 0.01. Peaks were annotated by their nearest genes using R function ChIPseeker::annotatePeak (*57*), with default dataset. The additional ChIP-seq data included to characterize the chromatin-associated factors is listed in Table S7 (*25, 26, 32, 42, 58-60*). We downloaded data sets from the ENCODE portal(*61*) (https://www.encodeproject.org/) with the following identifiers: ENCSR000ATQ, ENCSR000APX, ENCSR000AVB.

### PRO-seq data generation and analysis

We followed a previous published protocol (*62*) for PRO-seq. Briefly, around 5 million of hESCs were swelled in swelling buffer (10 mM Tris pH 7.5, 2 mM MgCl2, 3 mM CaCl2, 1XProteinase Inhibitor Cocktail, 10 U/ml Superase-In) for 10mins, then nuclei were extracted in swelling buffer added with 10% glycerol and 0.5% NP-40. The nuclei were resuspended in freezing buffer (50 mM Tris pH 8.3, 40% glycerol, 5 mM MgCl2, 0.1 mM EDTA, 1xPIC, 10U/100uL Superase-IN). Nuclear run-on (NRO) reaction takes place by mixing the nuclei in freezing buffer with equal volume of NRO buffer (10mM Tris pH 8.0, 5mM MgCl2, 300mM KCl, 1mM DTT, 100u/ml Superase.In, 1% sarkosyl, 250uM ATP and GRP, 50uM biotin-11-UTP and 50uM biotin-11-CTP) at 37°C for 5mins. The total RNAs from the nuclei were immediately extracted by TRIzol LS, and the NRO RNAs from this mixture were purified by streptavidin C1 beads twice. The purified NRO RNAs were processed to sequencing library using a two rounds of adaptor ligation following eCLIP-seq method with 10nt UMI (*63*). The libraries were sequenced using NextSeq 550 by the 40nt Pair-end model. We used Trim Galore (https://github.com/FelixKrueger/TrimGalore) to trim the adapters, and aligned all the clean reads to human reference genome hg19 using STAR 2.7.1(*64*). Then, we applied UMI-tools (*65*) to remove the duplicated reads. The unique aligned reads were further converted to bigwig format normalized by FPKM using BamCoverage function in deepTools (*66*).

### HOMER motif enrichment analysis

Motif enrichment on a set of genomic regions was performed by *HOMER v4.11.1* using the command *findMotifsGenome* (*31*). To compare one set of regions against another, we used the latter as background, instead of the random background regions *HOMER* uses by default.

### Comparative analysis of chromatin features signal

The ChIP-seq, ATAC-seq, or Pro-seq signal in a genomic region is estimated by the average score of all overlapping bins in the bigwig file, weighted by the width of their overlap with the given region. If there are no overlaps, we assume the score is zero. We compared the enrichment of signal in two groups of genomic regions by a *Wilcoxon rank-sum test*, also known as *Mann– Whitney U test* (*67*).

### Meta-peak analysis

The score matrix of a bigwig file in a set of genomic regions was calculated by *deepTools v3.5.1* (*68*), using the command *computeMatrix reference-point --referencePoint=center --binSize=1 -- averageTypeBins=mean*. To summarize the enrichment of signal in the regions, we calculated the mean score per base pair (i.e., average of each column in the score matrix).

### FOXA2 expression in hESCs

2x10^5^ iSAM hESCs were transduced with two lentiviral gRNAS targeting the promoter region of the endogenous *FOXA2* locus (see Table S9), aiming a MOI = 3, following the lentiviral infection protocol described above. After one day of recovery cells were cultured in E8 medium and treated with puromycin for three days. Dox was added to E8 media for three days and protein levels of GATA6 and FOXA2 were measured every 24 hours by flow cytometry analysis. To perform FOXA2 ChiP-seq experiments, iSAM hESCs transduced with the FOXA2 targeting lentiviral gRNAS were expanded for multiple passages in regular E8. Next, 2x10^7^ lenti-gRNA transduced cells were seeded in 15cm VTN-coated plates in E8 with Doxycycline and 24 hours later were collected to be crosslinked (see *ChIP-seq assays*).

### FOXA2 binding analyses based on motif enrichment

Genome-wide motif positions for human genome hg19 were downloaded from *HOMER* (*31*) (http://homer.ucsd.edu/homer/data/motifs/). Potential binding sites for FOXA1 and FOXA2 motifs in standard chromosomes (chr1, …, chr22, chrX, chrY) which were ≤20bp and annotated as “Intron” or “Distal Intergenic”, by *R* function *ChIPseeker::annotatePeak* (*57*), were selected and resized to 2Kbp around the center. Then, we filtered out regions that extended outside of chromosomes, overlapped with hg19 black-listed regions compiled by *ArchR* (*69*), or overlapped with FOXA2-bound regions (i.e., FOXA2 peaks detected upon activation at the ESC stage) which we also resized to 2Kbp around the center. From the top 5% of the remaining motif sites with the highest log odds scores, we created a non-overlapping set of regions by keeping the higher score out of each overlapping pair.

### Chromatin regions FOXA2 overlap and signal comparative analysis

Bed files for the chromatin regions were computed and subsets with any overlap were estimated using the package *rtracklayer* in R (*70*). ChIP-seq signal in the genomic regions of interest were estimated as the average score of all overlapping bins in the bigwig file.

### DNA EPIC-methylation analysis

The percentage of methylation in a genomic region is estimated by the average score of all overlapping bins in the corresponding bigwig file, weighted by the width of their overlap with the given region. If there are no overlaps, the level of methylation is unknown. Considering that the DNA methylation changes can extend for several hundred base pairs, we did not break down the differential ATAC peaks between the ESC and DE stages interrogated by the CRISPRa screen for *GATA6, EOMES, SOX17* and *MIXL1*. For this analysis, we redefined the CCRs by calling the ATAC peaks as competent if they overlapped a CCR (see *“Expanded CRISPRa screen analysis to identify CCRs for GATA6, EOMES, SOX17 and MIXL1 at the ESC stage”*) and as non-competent otherwise. For *PDX1* we used the same CCRs and non-CCRs previously defined. The final set of regions were extended by 500bp in each direction and used to compare the methylation levels.

### CRISPRa CCR targeting in QSER1/TET1-KO line

2x10^5^ *TET1/QSER1-KO* hESCs (DKO) and their corresponding WT matching line were transduced with the lentiviral supernatant generated from the Lenti-dCas9-VP64-Blast plasmid (addgene # 61425), after one day of recovery cells were selected under Blasticidin (10μg/ml) for 4 days. Next, cells were cultured seven days in E8 medium, and 1000 cells were seeded into VTN-coated 10cm plates. After 3 days of culture, 12 individual colonies were picked, grown for 5 days and the (d)Cas9 expression levels were checked by flow cytometry. DKO and WT colonies with comparable (d)Cas9 levels of expression were selected to be transduced with the lentiviral supernatant generated from a modified version of the Lenti-MS2-p65-HSF1-Hygro plasmid (addgene # 61426), where the Hygromycin cassette was replaced by the Zeocin selection cassette using In-Fusion cloning (Takara). After one day of recovery, cells were treated with Zeocin (conc) for 4 days. Next, cells were cultured seven days in E8 medium, 1000 cells were seeded into VTN-coated 10cm plates and after 3 days of culture, 5 individual colonies were picked, grown for 10 days and the MS2 and p65 expression levels were checked by qPCR. WT and DKO colonies with comparable expression levels were selected to perform CCR interrogation experiments. The resulting WT and DKO SAM hESCs lines were transduced with Lenti-Puro gRNAs targeting various validated CCRs from the studied genes, following the protocol described above. After a recovery day and 3 days of treatment with Puromycin (conc), cells were kept in E8 media for two additional days (in the same well) and fixed to measure the protein expression levels by flow cytometry analysis.

### Epigenetic screen

An aliquot of the Epigenetics Compound Library (Selleck L1900) was purchased from the MSK Gene Editing & Screening core. The library contains compounds including inhibitors of epigenetic enzymes such as Histone Deacetylase, SIRTs, Lysine demethylases, Histone Acetyltransferases, DNA Methyltransferase, and SIRTs activators (See table S8). The compounds were arrayed in 2 plates using a 96-well format as 1 mM stocks in DMSO (multiple wells containing DMSO only as controls). To perform the screen, two sets of 2x10^5^ iSAM hESCs were individually transduced with a lentiviral gRNA targeting the PDX1 CCR-1 and a non-targeting (NT) gRNA control. After transduction, cells were kept in E8 medium 2 days and treated with puromycin (conc) for 3 additional days and a final day of recovery in regular E8. 1x10^4^ cells from each group (PDX1 CCR and NT) were seeded into each well of four VTN-coated 96wp plates (total 8 plates), using E8 media with doxycycline (conc). The day after, fresh media was added and the compound in the library were transferred using a 96-pin replicator (V&P Scientific) at 1:1,000 (0.1% DMSO) to get 1μM final concentrations for each molecule. Media was changed 48 hours later with fresh E8 and doxycyline and a second round of the compound library was applied to each plate. Cells were kept two additional days, and PDX1-GFP expression levels were measured as described above using a BD LSR Fortessa and an Aurora cytometer (Cytek Biosciences) at MSKCC Flow Cytometry Core Facility.

### CCR Validation with HDAC inhibitors

iSAM-hESCs were transduced with representative lenti-gRNAs of the CCRs described in figure 6j and a non-targeting gRNA control as in “*Validation for individual CCRs by Flow Cytometry”*. After puromycin selection, cells were kept in regular E8 media and treated either with CI-994 (1μM) or DMSO for one day and for three additional days the treatment continued in presence of Doxycycline (2 μg/ml). At day 4, cells were fixed/stained and protein expression levels of the associated CCR genes were measured by flow cytometry.

### Schemes design

All the experimental schemes in this manuscript were created with BioRender (BioRender.com).

## Acknowledgments

We acknowledge the assistance from the following Memorial Sloan Kettering Cancer Center (MSKCC) Cores: Flow Cytometry, Gene Editing & Screening, Integrated Genomics Operation, Molecular Cytogenetics, and Stem Cell Research. We thank R. Garippa, and S. Mehta for assistance with CRISPR library generation and HiSeq for quantifying gRNA abundance after CRISPR screens, C. Cobbs for providing advice regarding the next-generation sequencing experiments, R. Gardner for assistance to design the sorting strategies for the CRISPR-based screenings, and S. Capellera for assistance with the scientific illustrations and writing of the manuscript.

## Author contributions

Conceptualization: JP, ZT, CSL, DH

Methodology: JP, ZT, WW, RL, SK, HC, JY, DM, AS, FG

Investigation: JP, ZT, DL, WW, RL, JRD, SK, JY, DM, RWR, AZ, FG, DY, WL

Visualization: JP, ZT, WW, RL, HC Funding acquisition: WL, TZ, EA, CSL, DH

Project administration: DH Supervision: JP, EA, CSL, DH

Writing – original draft: JP, DH Writing – review & editing: all authors

## Data and materials availability

Sequencing data are available at GEO under accession code <u>GSE233638</u>.

## List of Supplementary Materials

### Tables

**Table S1.** PDX1 CRISPRa and CRISPRi screening results to discover PDX1

**Table S2.** PDX1 CCRs validated and discovered

**Table S3.** CRISPRa extended screening results to discover GATA6, SOX17, EOMES and MIXL1 CCRs

**Table S4.** TADs called from Hi-C assays in hESCs

**Table S5.** 4C viewpoint oligos

**Table S6.** ChIP-seq data sets used to characterize the chromatin-associated factors enriched in CCRs

**Table S7.** HOMER motif enrichment analyses on CCRs

**Table S8** Epigenetic compound library description

**Table S9.** List of oligos used (including gRNAs)

**Table S10.** List of antibodies used

---

**Supplementary Figure 1.**
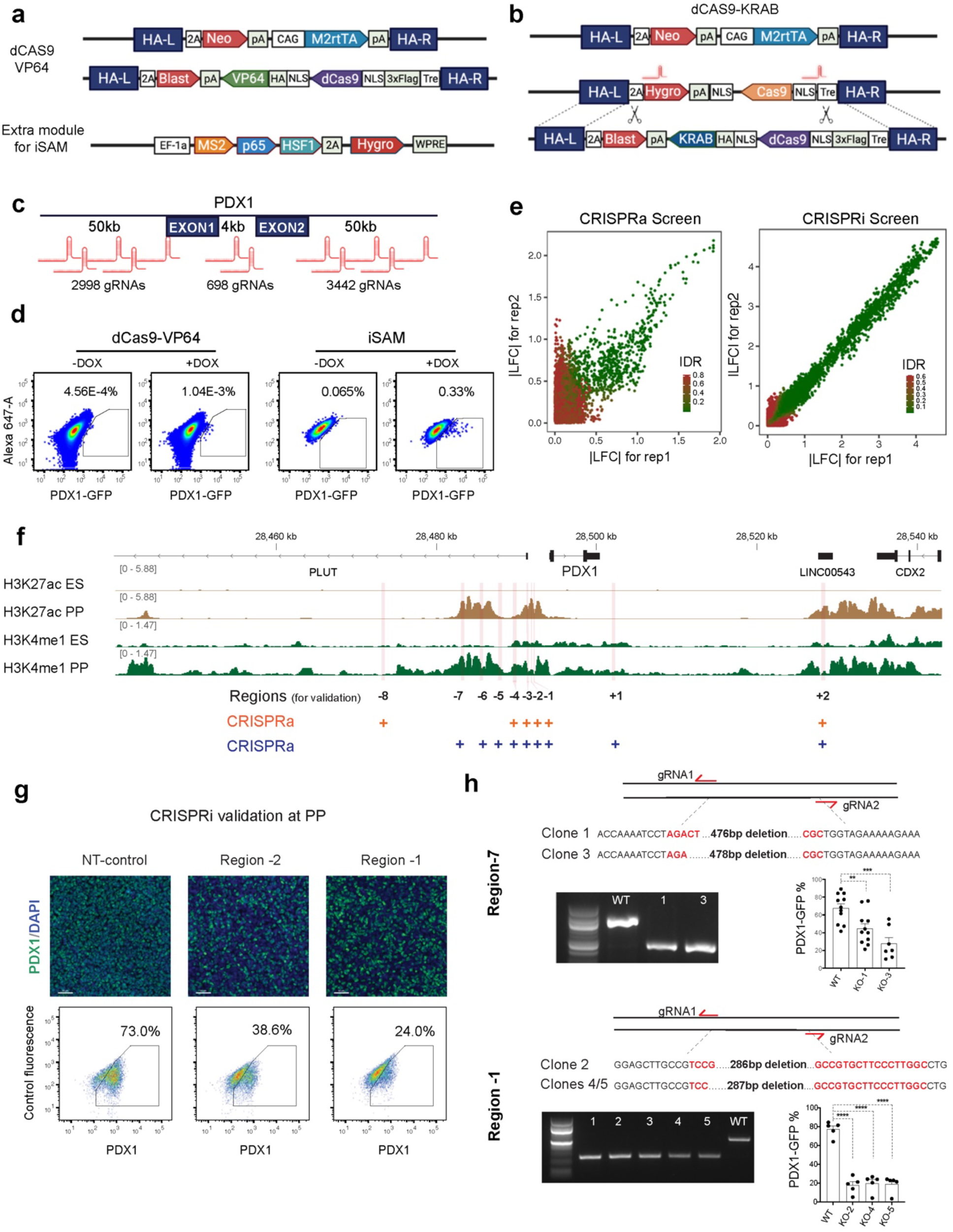
**a)** Insertion cassettes targeted to the AAVS1 locus to derivate an inducible dCas9-VP64 hESC line starting from a previously designed iCas9 H1 ESC line. To generate the iSAM hESC line, the dCas9-VP64 line was transduced with an additional lentivirus (lower panel). **b**) Cloning strategy to generate the inducible dCas9-KRAB line using as a background the iCas9 H1 ESC line. **c)** Schematics of the full-tiled gRNA library designed to interrogate the presence of cis-regulatory regions that control the expression of PDX1. **d)** Pilot experiments showing the presence of PDX1+ cells at the ESC stage after transducing the dCas9-VP64 hESC and the SAM-hESC line with the tiled PDX1 gRNA library. After transduction with the lenti-gRNA library, cells were treated with doxycycline to activate the expression of the CRISPRa system (see methods). **e)** Reproducibility between replicates of the screens performed with the SAM-hESC line at the ESC stage (left panel) and the dCas9-KRAB line (right panel) during the differentiation from ESC to the PP stage (IDR was calculated between the z-scores of the LFC values). **f)** ChIP tracks of histone marks previously associated to enhancer regions around the PDX1 region at the ES and PP stages (pink bars: candidate regulatory regions from the screens). **g)** Immunostaining images and FACS dot plots showing PDX1 expression levels after interrogation of a subset of candidate regulatory regions with the dCas9-KRAB tool during the differentiation from ESC to PP. A non-targeting gRNA was used as a control (see Methods) (PDX1 = Green, DAPI = blue). **h)** Validation of regions -7 and -1 as PDX1 enhancers by targeted-deletion experiments. The targeting strategy to remove the PDX1 CCR -7 (left panel) and CCR -1 (right panel) using an iCas9-hESC line is described (see methods). Sequencing results of the individual clones validated for each CCR and box plots summarizing the percentage of PDX1+ cells obtained after differentiation to the PP stage for each deletion line compared to the WT line are shown.

**Supplementary Figure 2.**
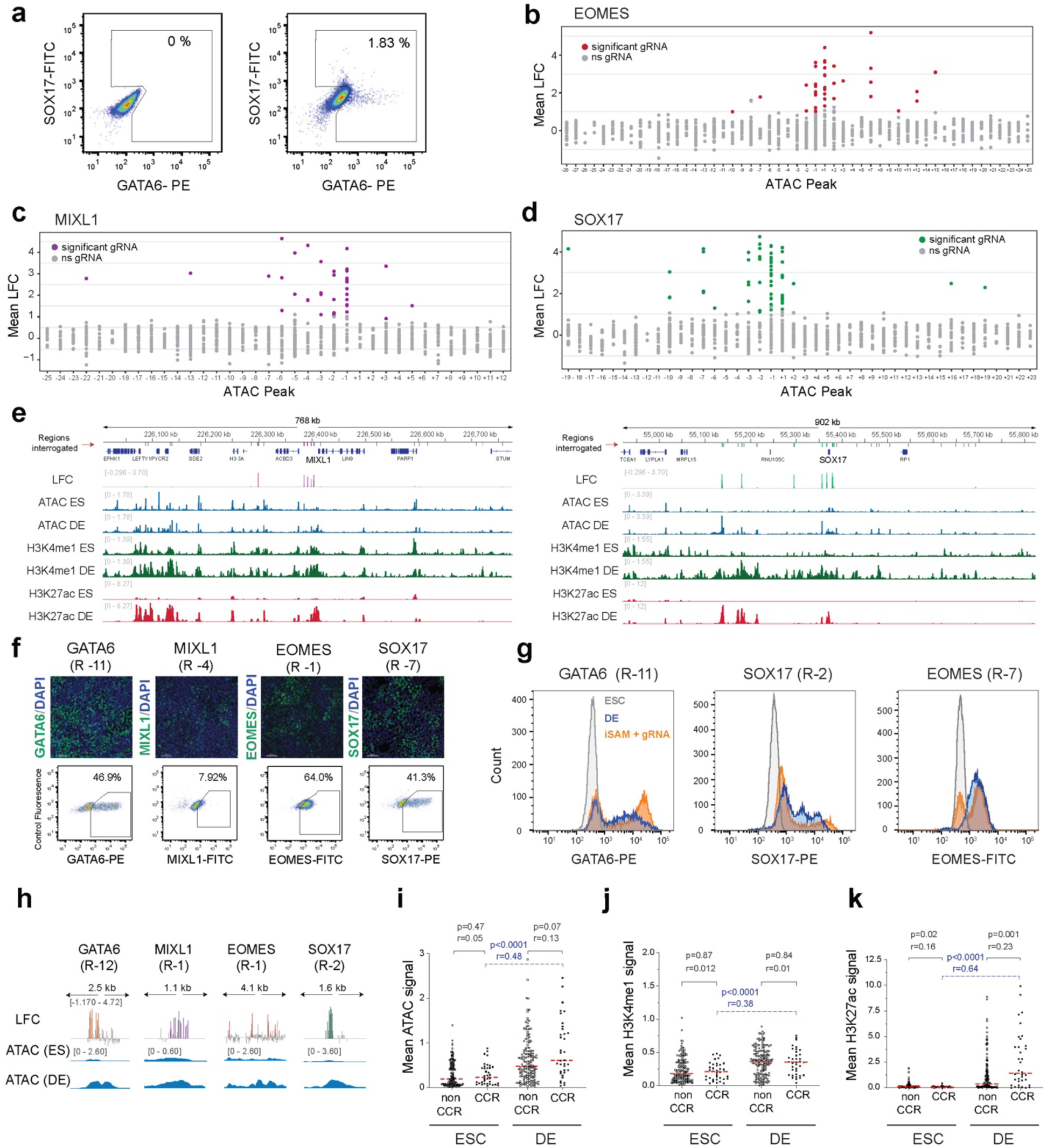
**a**) FACS dot plot to illustrate the sorting strategy to collect any GATA6 or SOX17+ cells after transduction of the iSAM hESC line with a customized gRNA library targeting candidate CCRs. **b-d)** Mean LFC distribution of the gRNAs present in the positive fraction of the sorted cells after CRISPRa interrogation. Each dot is a gRNA color-coded as in Fig. 2b (ns= non-significant) plotted across the interrogated regions surrounding EOMES(n=53), MIXL1(n=37) and SOX17(n=42). **e)** Multi-track plot to illustrate the signal distribution of features commonly associated to active or primed enhancers in hESC and DE cells at the interrogated regions around the MIXL1 and SOX17 loci (Interrogated regions and each bar in the LFC track are color coded as in Fig. 2b). **f)** Representative immunofluorescence images (*upper panel*) and FACS plots (*lower panel*) to show the effect of activating individual CCRs via gRNA-targeting with the SAM tool for each gene (scale bar =80um). **f)** Representative histograms comparing the protein expression levels for GATA6, SOX17 and EOMES in ESCs (gray), differentiated Definitive Endoderm cells (blue) and iSAM hESCs targeting a CCR (indicated on top) associated to each gene (orange). **h)** High magnification of the gRNA distribution along four representative CCRs for each interrogated gene (color-coded as in 2b). **i-k)** ATAC, H3K4me1 and H3K27ac average signal in CCRs and non-CCRs at ESC and DE stage (red dotted line = median) (Wilcoxon rank-sum test p values and effect sizes (r) are included).

**Supplementary Figure 3.**
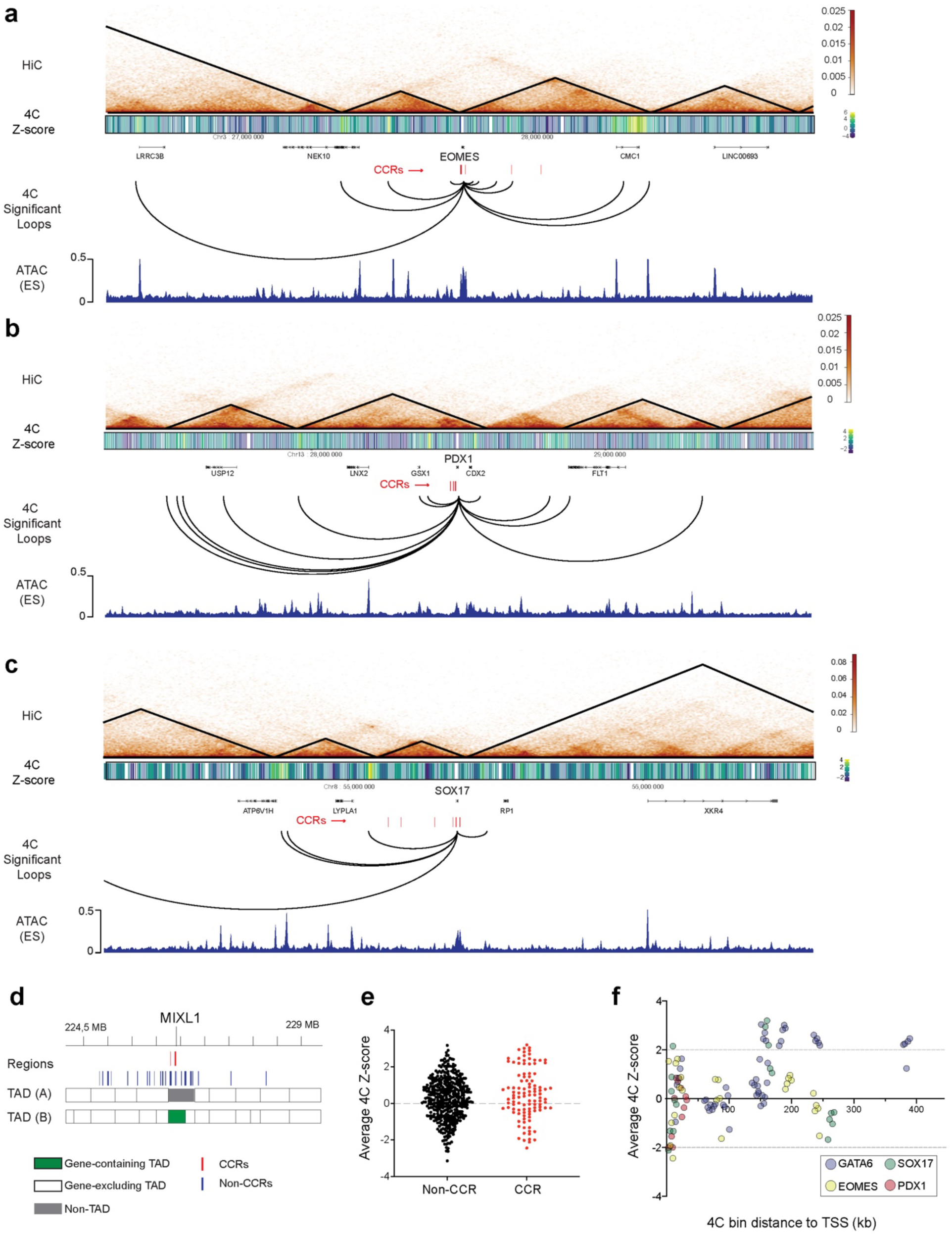
**a-c)** KR normalized Hi-C plots around EOMES, PDX1 and SOX17. Heatmap of the 4C Z-scores using as viewpoints the gene bodies of each gene and 4C significant contacts (q ≤ 0.1) are depicted. ATAC tracks at the ES stage are included for reference. **d)** TAD surrounding the MIXL1 gene locus discovered by HiC assays in hESC, two TAD-calling strategies are compared (A=5 bins, B=10 bins, see Table S4 and methods). **e)** Z-score value for the 4C bins overlapping with at least one CCR or a non-CCR across the four interrogated genes. **f)** Z-score values of the 4C bins overlapping with at least one CCR across the genomic locations of the four interrogated genes taking as a reference the Transcription Start Site (TSS) of each gene.

**Supplementary Figure 4.**
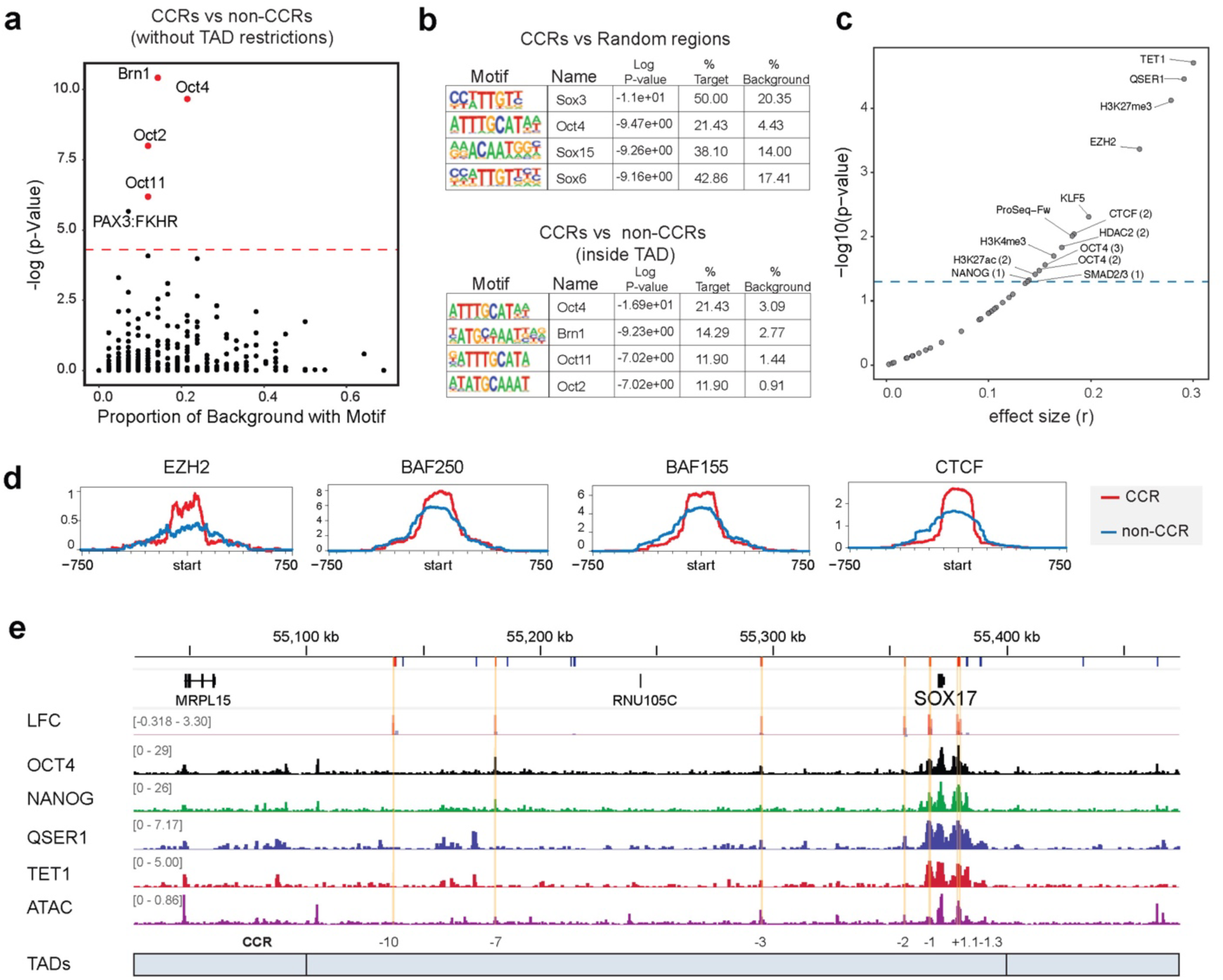
**a)** HOMER motif enrichment analysis plot to summarize the significantly enriched motifs (above the red-dotted line) when comparing CCRs vs all non-CCRs. **b**) Description of the top enriched motifs in the CCRs discovered by HOMER, compared to random (above) and non-competent regions inside the TADs containing the interrogated genes (below). **c)** ChIPseq data was used to compare the signal enrichment at the ESC stage in the CCRs vs all non-CCRs (See methods). Factors that bind significantly differently between CCRs and non-CCRs are indicated above the blue-dotted line (Wilcoxon rank-sum test, p< 0.05). **d)** Average ChIP-seq scores of chromatin features measured at CCRs and non-CCRs in hESCs (Wilcoxon rank-sum test, p values and effect sizes indicated on top). **e)** Average of normalized counts visualized as a MetaPeak using multiple ChIP-signal data sets in CCRs (red) compared to non-CCRs(blue). **f)** Multi-track plot to illustrate the ChIPseq signal of the enriched factors at the interrogated regions around the SOX17 locus, including TADs (Fig 4f zoomed out). Interrogated regions are depicted in red boxes, and competent regions are highlighted in yellow. (LFC = log Fold change in the CRISPRa screen).

**Supplementary Figure 5.**
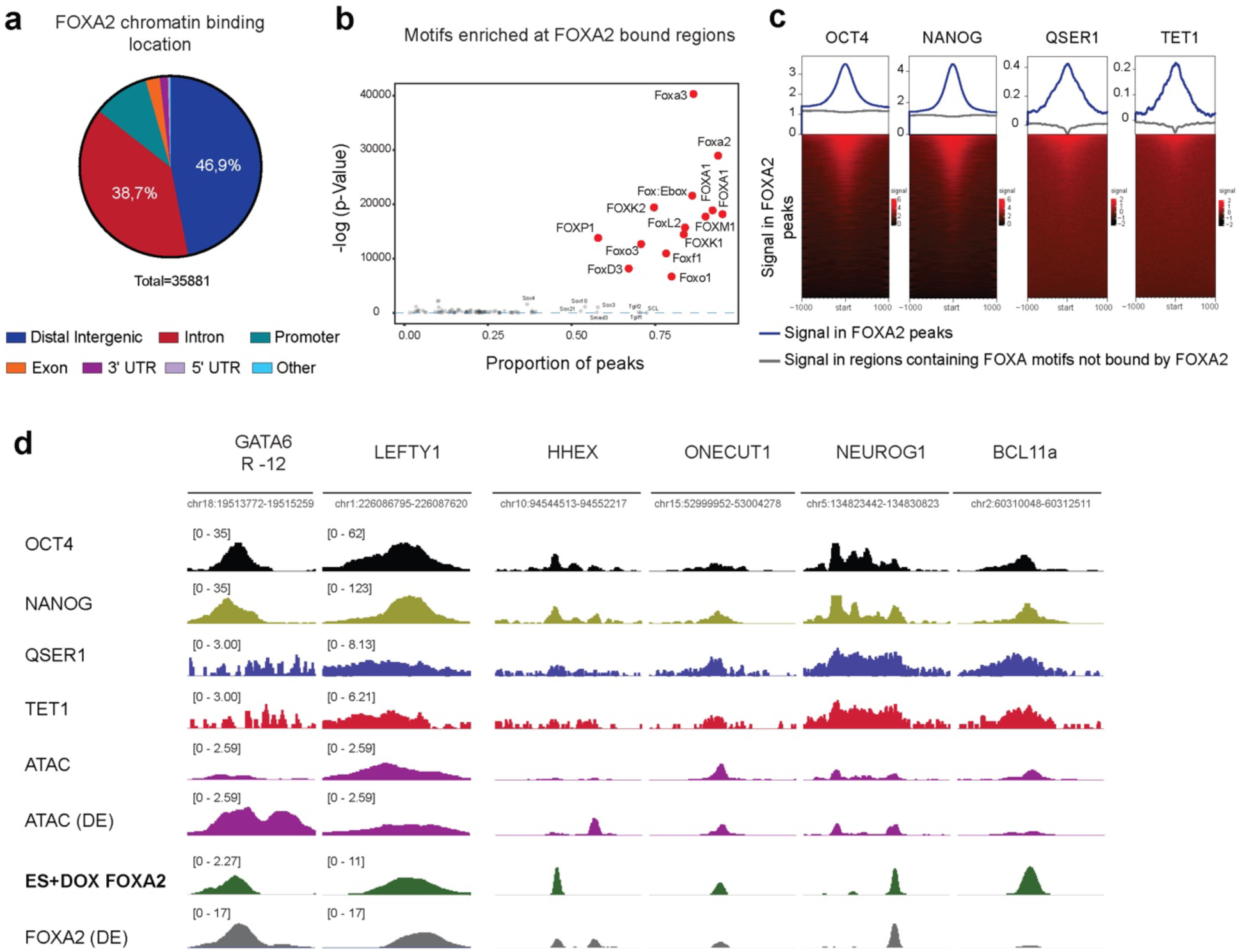
**a)** FOXA2 binding genomic distribution in hESCs. **b)** HOMER motif analysis plot to summarize the significantly enriched motifs (above the blue-dotted line) in the FOXA2 peaks detected upon activation at the ESC stage (including FOX specific motifs). **c)** Average of normalized counts visualized as meta-peaks to illustrate ONQT ChIP-signal at the regions bound by FOXA2 in hESCs compared to the signal at equivalent regions enriched in FOXA motifs not bound by FOXA2. **d)** Multi-track plot of regions containing FOXA2 peaks upon its activation at the ESC stage and the ONQT factors at GATA6 CCR -12 and various putative regulatory regions of the genes indicated (upper label). For comparison purposes ATAC peaks and FOXA2 peaks found at the Definitive Endoderm stage (DE) are included.

**Supplementary Figure 6.**
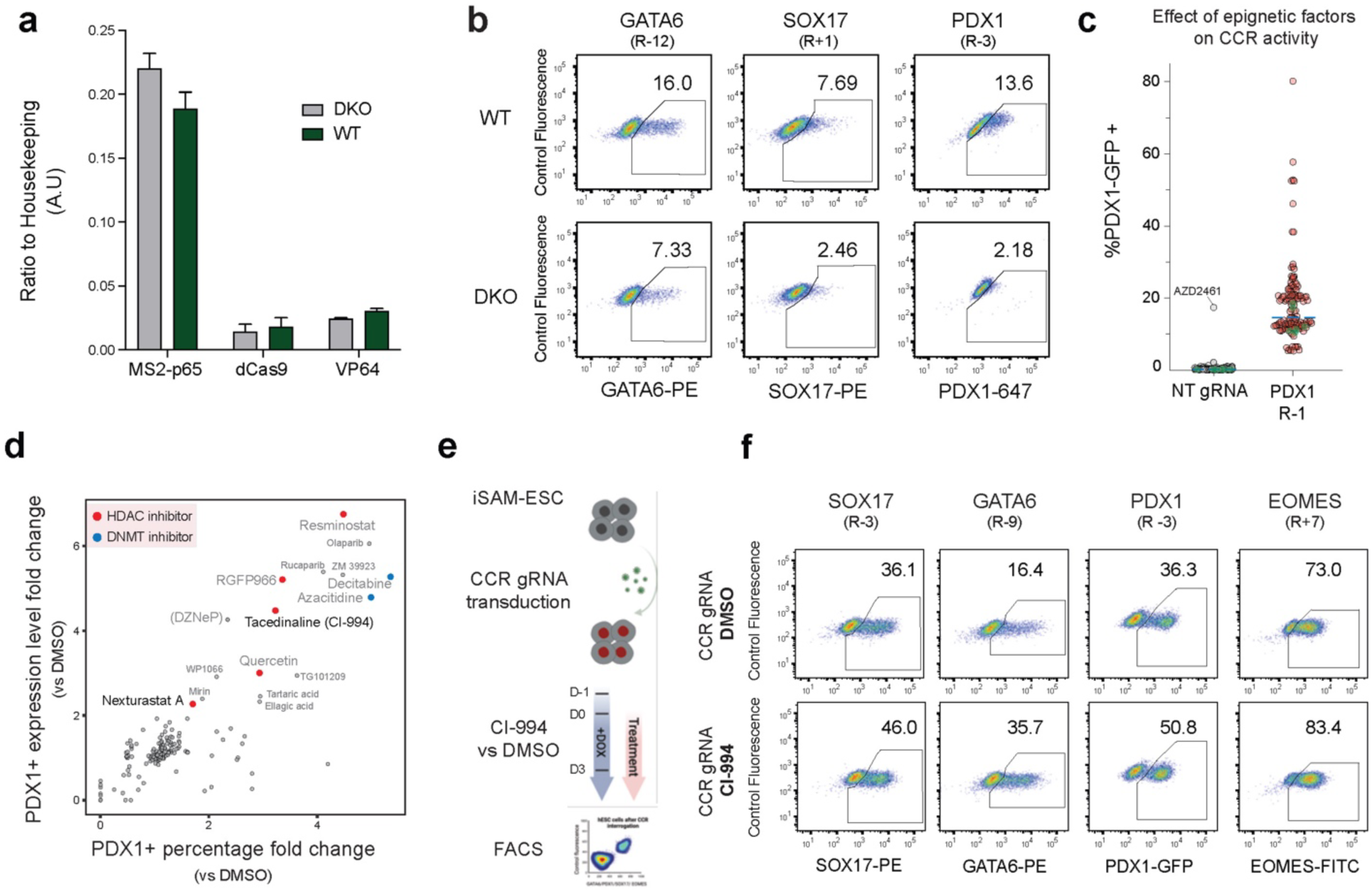
**a)** Quantitative PCR expression analyses of the exogenous genes associated to the SAM activation system in WT and DKO hESC lines (n>2). **b)** Representative FACS dot plots to illustrate the protein expression levels associated to the interrogated genes after activation of several CCRs in WT and DKO hESCs (gene and the associated CCR are indicated on top). **c)** Percentage of PDX1 positive cells after treating hESCs with an epigenetic library during the CRISPRa interrogation of the PDX1 CCR-1 vs a non-targeting (NT) gRNA control. Each dot corresponds to a single molecule of the library (Green dots = DMSO, blue-dotted line = median). **d)** Scatter plot that summarizes the results of the epigenetic screen. Each dot corresponds to the average fold change of the percentage of PDX1 positive cells and the PDX1 protein levels after interrogation of the PDX1 CCR-1 and treatment with each molecule of the library, compared to the DMSO controls (See methods). The top categories of epigenetic modulators that significantly increase PDX1+ levels are color-coded. (Grey fonts to indicate compounds that affect cell proliferation at the employed dose). **e)** Experimental design to test the effect of the HDAC1-family inhibitor CI-994 in the transcriptional competence of multiple CCRs. **f)** Representative FACS dot plots to illustrate the effect of the CI-994 treatment on the transcriptional competence of a CCR after interrogation by the SAM system. *Upper panels*: cells transduced with a gRNA targeting a CCR of the gene annotated above and treated with DMSO. *Lower panel*: cells transduced with a gRNA targeting a CCR of the gene annotated above and treated with CI-994.

